# Photosymbiotic algae acquisition and their interactions with the acoel *Convolutriloba macropyga*

**DOI:** 10.64898/2026.03.06.710013

**Authors:** Francesca Pinton, Gloria Lando, Viviana Cetrangolo, Katja Felbel, Elisabeth Grimmer, Andreas Hejnol, Nadezhda N. Rimskaya-Korsakova

## Abstract

Symbiosis with photosynthetic microbes is widespread in marine animals, with various symbiont transmission modes and localisation within the host. Here, we characterise the association between the acoel *Convolutriloba macropyga* and its photosymbionts, identified as *Tetraselmis* green algae based on rbcL gene phylogenetic analysis. Symbionts are transmitted vertically to asexual offspring and acquired horizontally by juveniles after sexual reproduction. Embryos develop to aposymbiotic juveniles that ingest *Tetraselmis* through the mouth. Confocal microscopy shows an increase in algae number within juveniles and in their presence at the body wall. Transmission electron microscopy reveals that symbionts lose flagella and theca. In adults, symbionts are extracellular at the body periphery, but can be intracellular within the parenchyma, in contrast with previously described acoel photosymbionts. This likely reflects different host-symbiont interactions, with algae potentially performing photosynthesis and nutrient exchange at the periphery, while undergoing transport or digestion in the parenchyma. Comparative transcriptomics between symbiotic adults and aposymbiotic juveniles shows an enrichment of amino acid synthesis, lipid metabolism, and osmotic and oxidative stress responses in symbiotic adults. Our data shows that algal symbionts engage with host tissues in distinct ways, inside or outside host cells, highlighting a previously unappreciated spatial complexity in host–algae interactions.

**Highlights:**

- Tetraselmis algae are taken up by Convolutriloba macropyga juveniles
- Algal symbionts in juveniles lose theca and flagella, proliferate, and move to the body wall
- Symbionts are extracellular at the body wall and can be intracellular in the parenchyma
- Amino acid synthesis, lipid metabolism, osmoregulation and stress responses are activated in symbiotic adults

## Introduction

Symbioses between microbes and animal hosts are ubiquitous. Beside influencing their hosts’ metabolism and development, symbionts shape their ecology and evolution (Bosch & McFall-Ngai, 2011). Effects of the symbionts on the host can range from advantageous to detrimental, but many provide benefits such as nutrition, waste recycling, protection from pathogens, or even bioluminescence (Chomicki et al., 2020; Davy et al., 2012). In some associations, endosymbionts are passed on directly to the next generation (vertical transmission), whereas in others, an aposymbiotic (i.e. non-symbiotic) organism must acquire symbionts from the environment (horizontal transmission). In vertical transmission, symbionts are often associated with the host throughout its entire life cycle, with no aposymbiotic phase. Therefore, vertical transmission needs a potentially complex strategy to transfer the symbionts to the next generation, but it also enables a more specialised and fine-tuned relationship (Bright & Bulgheresi, 2010). Horizontal transmission allows for more flexibility, but it requires mechanisms to recognize the symbiont and establish the partnership, as well as to maintain it over time (Bright & Bulgheresi, 2010; Davy et al., 2012). The transmission mode and the interactions between host and symbionts affect their co-evolution (Bright & Bulgheresi, 2010). They are inherently affected by the location of the symbionts (Bright & Bulgheresi, 2010; Chomicki et al., 2020), which are often confined to a specific cell type, tissue, or organ (Dubilier et al., 2008; Fronk & Sachs, 2022; Liao et al., 2025).

One specific kind of symbiosis, photosymbiosis, entails a heterotrophic host and photoautotrophic endosymbionts (Liao et al., 2025). This type of symbiosis—usually considered mutualistic, i.e. beneficial for both partners—has evolved in many animal hosts, from sponges and cnidarians to ascidians and molluscs, which can associate with unicellular “algae”: cyanobacteria, chlorophytes, dinoflagellates, diatoms, or even individual chloroplasts (Melo Clavijo et al., 2018). The symbionts provide the host with O_2_ and photosynthesis products such as sugars and lipids, and they receive protection, CO_2_, and inorganic nutrients, mainly nitrogen compounds (Davy et al., 2012; Liao et al., 2025; Melo Clavijo et al., 2018; Yee et al., 2025). In the best studied example of animal photosymbiosis, the cnidarian-dinoflagellate association, the endosymbionts can be either vertically transmitted or horizontally acquired by aposymbiotic polyps or larvae (Bucher et al., 2016; Muller-Parker et al., 2015). They are found in cells of the gastrodermis within symbiosomes: complexes of membranes kept at an acidic pH, which favours photosynthesis (Davy et al., 2012; Liao et al., 2025). The symbionts are enclosed in a symbiosome in other systems, too: chlorophytes in the freshwater green hydra, chlorophytes or dinoflagellates in sponges (Liao et al., 2025), and chlorophytes in spotted salamanders (Kerney et al., 2011). On the other hand, symbiotic cyanobacteria can be either intra- or extracellular, in the mesohyl of sponges and in the tunic or cloaca of ascidians (Hirose, 2015; Mutalipassi et al., 2021). In giant clams, extracellular dinoflagellate symbionts are confined in specialized mantle tubules (Liao et al., 2025).

Acoela are soft-bodied motile marine bilaterians. Some species can host **dinoflagellates as photosymbionts intracellularly** (Barneah et al., 2007; Hikosaka-Katayama et al., 2012; Ogunlana et al., 2005; Zhong et al., 2025), but other species host **chlorophytes as extracellular symbionts** (Bailly et al., 2014). The hosts cannot survive without algae, i.e. the symbiosis is **obligate** (Åkesson & Hendelberg, 1989; Apelt, 1969; Arboleda et al., 2018; Shannon & Achatz, 2007). The transmission mode is horizontal in *Symsagittifera roscoffensis, Praesagittifera naikaiensis,* and *Convoluta convoluta* (Bailly et al., 2014; Hikosaka-Katayama et al., 2024; Pedersen, 1964), but maternal transmission has been described for dinoflagellates in *Waminoa* (Barneah et al., 2007; Hikosaka-Katayama et al., 2012). *Convolutriloba,* on the contrary of other photosymbiotic acoels, can reproduce both **sexually and asexually**, with ***Convolutriloba macropyga* being the only species with a prolific and regular sexual reproduction** (Shannon & Achatz, 2007) on top of asexual reproduction by budding.

Algal symbionts seem not to be confined to specific acoel organs or tissues, but this lacks in depth research; even their extracellularity still requires confirmation (Hirose & Hirose, 2007; Zhong et al., 2025). The uptake in time and space after reproduction is a question only partially addressed, as well as how it is affected by symbiont transmission mode (Barneah et al., 2007; Douglas, 1983; Hikosaka-Katayama et al., 2012; Oschman, 1966). Moreover, our understanding of the interactions of the endosymbionts with host tissues and cells remains very limited.

## Results

### *C. macropyga* embryos develop aposymbiotic, juveniles acquire *Tetraselmis* algae from the environment

Observations of budding adults leave no doubt that green algal symbionts in *Convolutriloba* sp. can be transmitted vertically during asexual reproduction (Fig. 1A) (Åkesson et al., 2002; Shannon & Achatz, 2007). When the animals lay eggs, however, juveniles contain no symbionts upon hatching—i.e. they are aposymbiotic— as described in (Shannon & Achatz, 2007) and confirmed by our documentation of embryo development (Fig. 1B). The sexually produced progeny must therefore acquire symbiotic algae horizontally from the environment and establish photosymbiosis from scratch at each generation. We describe the development of *Convolutriloba macropyga* and experimentally characterise the uptake of *Tetraselmis* algae by their juveniles.

**Fig. 1.**
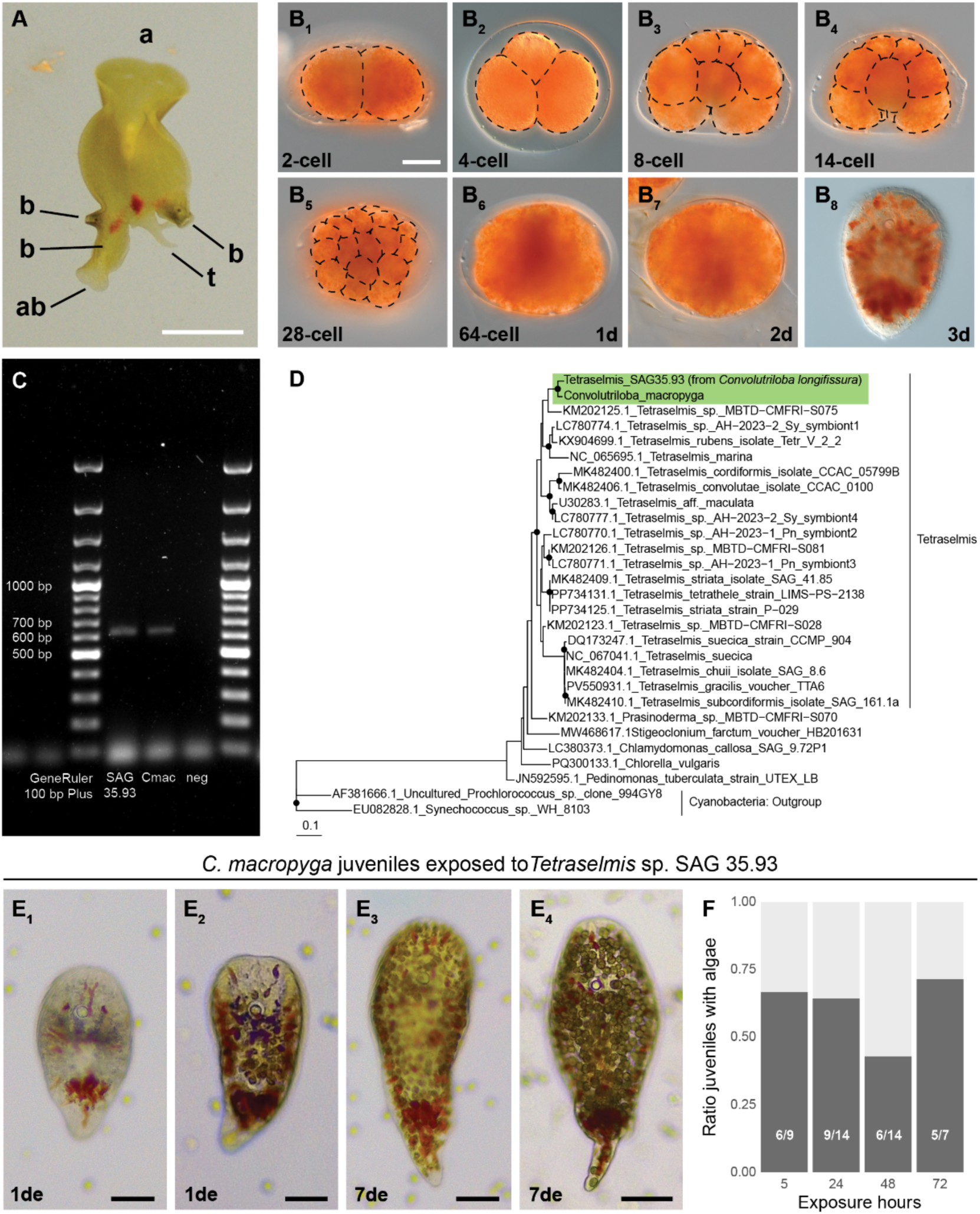
Development of *C. macropyga* and uptake of *Tetraselmis* sp. Algae. **A** Adult *C. macropyga* undergoing asexual reproduction. Three individuals budding from the posterior side of the animal (b), at various stages of development. Dorsal view, with anterior side facing up (a); note that polarity of the asexual offspring is reversed (ab: anterior of budding individual); t: tail. **B_1_- B_8_** Embryonic development of *C. macropyga.* **C** Gel electrophoresis of PCR products obtained with rbcL primers and cDNA from free living *Tetraselmis* sp. SAG 35.39 (SAG 35.39), cDNA from *C. macropyga* adults (Cmac), and nuclease-free water as a negative control (neg). **D** Phylogenetic tree of partial rbcL, i.e. sequenced PCR products from free living *Tetraselmis* sp. SAG 35.39 and *C. macropyga* adults (highlighted in green), together with sequences from other *Tetraselmis* sp., chlorophytes, and cyanobacteria as an outgroup; nodes with a black dot have Ultrafast Bootstrap support value > 90; scalebar is number of expected substitutions per site. **E_1_-E_4_** Morphological changes of *C. macropyga* juveniles and uptake of *Tetraselmis* algae: **E_1_** 1 day of *Tetraselmis* exposure (de), no algae inside the animal, rounder shape; **E_2_** 1 de, few algae in the central part of the body, longer shape; **E_3_** 7 de, algae in the whole body, longer shape, tail thinning; **E_4_** 7 de, algae in the whole body, thinner tail. **F** Ratio of juveniles containing algae after exposure to *Tetraselmis* for different time windows (Pearson’s Chi-squared test, χ_2_ = 2.337 p = 0.51); the actual ratios are shown on the bars. Lateral view, animal side up in **B_1_-B_7_**; dorsal view, anterior side up in **B_8_,E**. Scale bar: 2 mm in **A**, 50 µm in **B,E.**

Embryonic development from egg-laying to hatching of the juveniles lasts 3 days (Fig. 1B). The embryo follows the typical acoel duet cleavage: the first division is equal and follows the animal-vegetal axis (Fig. 1B_1_). The second cleavage, perpendicular to the animal-vegetal axis, gives rise to two smaller cells at the animal pole (micromeres) and two bigger cells at the vegetal pole (macromeres): the first micromere and macromere duets (Fig. 1B_2_). After a cleavage of the macromeres followed by a cleavage of the micromeres, the embryo reaches the 8-cell stage (Fig. 1B_3_). Gastrulation begins at 14-cell stage by ingression of the macromeres at the vegetal pole and lasts approximately 50 minutes (Fig. 1B_4_). During this process, micromeres at the animal pole continue to divide and by the end of gastrulation, the embryo is formed by 28 cells (Fig. 1B_5_). Division of ectodermal micromeres results into a layer of ciliated epidermal cells (Fig. 1B_6_). Afterwards, organs and tissues form, while the embryo rotates inside the eggshell, until the newly formed juvenile hatches at around 3 days from egg-laying (Fig. 1B_7_-1B_8_). The drop-shaped morphology of hatchlings differs greatly from the morphology of adults and older juveniles containing algae (Fig. 1A,E).

Within 5 days from hatching, *C. macropyga* juveniles were exposed to potential symbionts (*Tetraselmis* sp. SAG 5.93 isolated from *Convolutriloba longifissura* (Balzer, 1999; Bartolomaeus & Balzer, 1997)). The close relatedness of SAG 35.93 and *C. macropyga* symbionts is supported by the phylogenetic analysis of rbcL (Rubisco; Ribulose-1,5-bisphosphate carboxylase/oxygenase), amplified from adult *C. macropyga* holobiont and free-living *Tetraselmis* sp. SAG 35.93 (Fig. 1C,D, Data S1,S2). These results also confirm that *C. macropyga* and *C. longifissura* symbionts are closely related, and they belong to the *Tetraselmis* clade of Chlorophyta (Fig. 1D).

When *Tetraselmis* are available in the medium, juveniles feed on them (Video S1). Distinct feeding-related behaviours can be observed: (1) flow of medium and algae towards the animal, likely due to cilia movement (Video S1, S2); (2) the anterior half of the body forms a funnel, as observed in *Convolutriloba* adults (Bartolomaeus & Balzer, 1997; Shannon & Achatz, 2007) (Video S1); (3) swimming in circles leads to the amassing of algae on the ventral side, where the mouth is (Video S2, S3). The algae can be observed in the whole body of some *C. macropyga* juveniles, which become more elongated and with thinner tails over time (Fig. 1E). Only some juveniles take up *Tetraselmis* algae, independently of exposure time (Fig. 1F).

### The number of algae within juveniles and in association with their body wall increases over time

Algae autofluorescence—coupled with DAPI and phalloidin, staining host nuclei and muscle fibres, respectively—enabled us to characterise the uptake of algae within *C. macropyga* juveniles (Fig. 2A). We expected cell proliferation and a gradual migration from the parenchyma surrounding the mouth towards the body periphery, where the sunlight is more easily available. Specifically, we quantified the number of algae acquired and their localization within the juveniles up to 5 days from the start of exposure. Juveniles were exposed to algae for intervals ranging from 5 hours to 3 days (exposure time, ET); then algae were removed for varying amounts of time (Post-Exposure Days, PED). We also assess changes occurring across the Days from Exposure Start (DES), encompassing both exposure and post-exposure time. Algal abundance within the juveniles increases significantly with exposure time (Fig. S1A), with DES (Fig. 2B), and with PED (Fig. 2C), suggesting continuous *Tetraselmis* uptake by *C. macropyga* juveniles and proliferation of *Tetraselmis* within juvenile tissues.

**Fig. 2.**
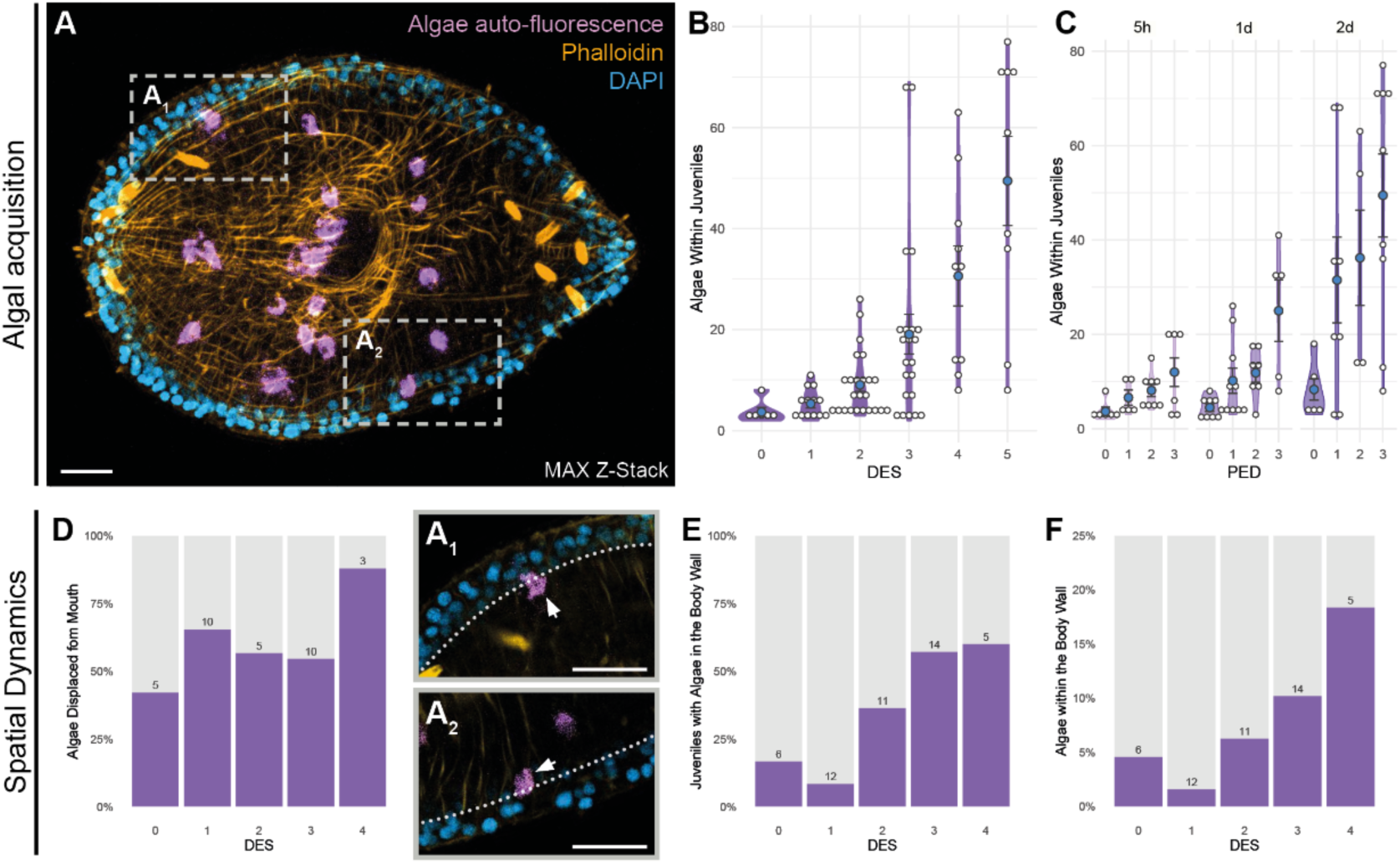
*Tetraselmis* sp. abundance and spatial distribution within *C. macropyga* juveniles. **A** Maximum projection of a confocal image stack of *C. macropyga* juvenile (ET 2d, PED 0). Blue: DAPI; yellow: phalloidin; purple: algae autofluorescence. **A_1_ – A_2_** Single focus planes showing details of body wall; dotted lines highlight the boundary of the body wall. Algae classified as body-wall-associated are marked with white arrowheads. **B – C** Number of algae present within juveniles, minimal adequate model: log(Algae_number) ∼ time, **B** time is DES (Adj. R^2^=0.461, p=2.123e-10), **C** time is PED, grouped by exposure time (5h ET: Adj. R^2^=0.217, p=0.039; 1d ET: Adj. R^2^=0.390 p=1.332e-4; 2d ET: Adj. R^2^=0.272, p=0.014). **D** Percentage of algae located far from the mouth relative to z axis (Pearson’s Chi-squared test, χ^2^=12.758, p=0.012); sample size above columns. **E** Percentage of juveniles with algae associated with body wall (Pearson’s Chi-squared test, χ^2^=8.9849, p=0.06148); sample size above columns. **F** Percentage of algae associated with body-wall (Pearson’s Chi-squared test, χ^2^=14.843, p=5.038e-3); sample size above columns. Scale bars are 20 µm.

We then quantified algal displacement from the mouth opening. The position on the xy plane does not change with post-exposure time (Fig. S1B), even when normalized to juvenile body size (Fig. S1C). In contrast, the number of algae far from the mouth on the z axis differs with DES (Fig. 2D, Fig. S1D), indicating a change in position along the dorso-ventral axis. Additionally, algae are displaced mainly towards the dorsal side (Fig. S1E).

We further examined the presence of algae in the body walls, as an additional indicator of movement towards the body periphery. Juvenile nuclei stained with DAPI were used to delineate the limits of the body wall (Fig. 2A,A_1_,A_2_). The number of juveniles containing body-wall-associated algae increases, but not significantly, with DES and PED (Fig. 2E, Fig. S1F). The number of body-wall-associated algae varies significantly with DES (Fig. 2F), suggesting the association of algae with the body wall within a few days from uptake.

We also investigated symbiont dynamics in adults, by providing them with *Tetraselmis* sp. SAG 35.93 (potential symbionts) or brine shrimps (food) regularly for 19 days. In both samples, the ratio between algae and animal cells is comparable to the one in controls (Fig. S2A), while animal size is significantly, but slightly, larger when food is available (Fig. S2B). Therefore, adult *C. macropyga* are unlikely to continuously take algae up from the environment, because the algae-animal cell ratio is not higher when algae are available in the medium; however the number of algae is likely to increase by proliferation, if food is available, because the algae-animal cell ratio is not affected by food availability but animal size is.

### *Tetraselmis* algae lose theca and flagella upon uptake by *C. macropyga* juveniles

We then looked for ultrastructural changes in *Tetraselmis* algae upon uptake by *C. macropyga* juveniles. Free living *Tetraselmis* sp. algae used for symbiosis establishment experiments are always observed in a non-swimming state (Video S4). Their ultrastructural morphology resembles other *Tetraselmis* sp. algae (Arora et al., 2013; Hori et al., 1982; Hyung et al., 2021; Manton & Parke, 1965; Norris et al., 1980; Parke & Manton, 1967; Riewluang & Wakeman, 2025). Although 4 short flagella emerge from a cell depression—the flagellar pit (Norris et al., 1980)—, they are completely contained within the theca (Fig. 3A,D,E,F). We also observe some cells without flagella and with a rounder morphology and/or multiple theca layers (Fig. 3B,C), which can be interpreted as encysting cells (Kang et al., 2024; Norris et al., 1980; Parke & Manton, 1967). TEM reveals the presence of an eyespot (Fig. 3G) and a two-layer theca (Fig. 3DG): the inner theca is a 55nm-wide electron dense layer, the outer one is 35 nm wide with hairs towards the outside (Fig. 3G). In some instances, two algal cells share an outer theca, while retaining their inner theca (Fig. 3H,I): we interpret them as sister cells after cell division. We also observe a loss of structural integrity in some algal cells, with the inner theca is lost (Fig. 3J).

**Fig. 3.**
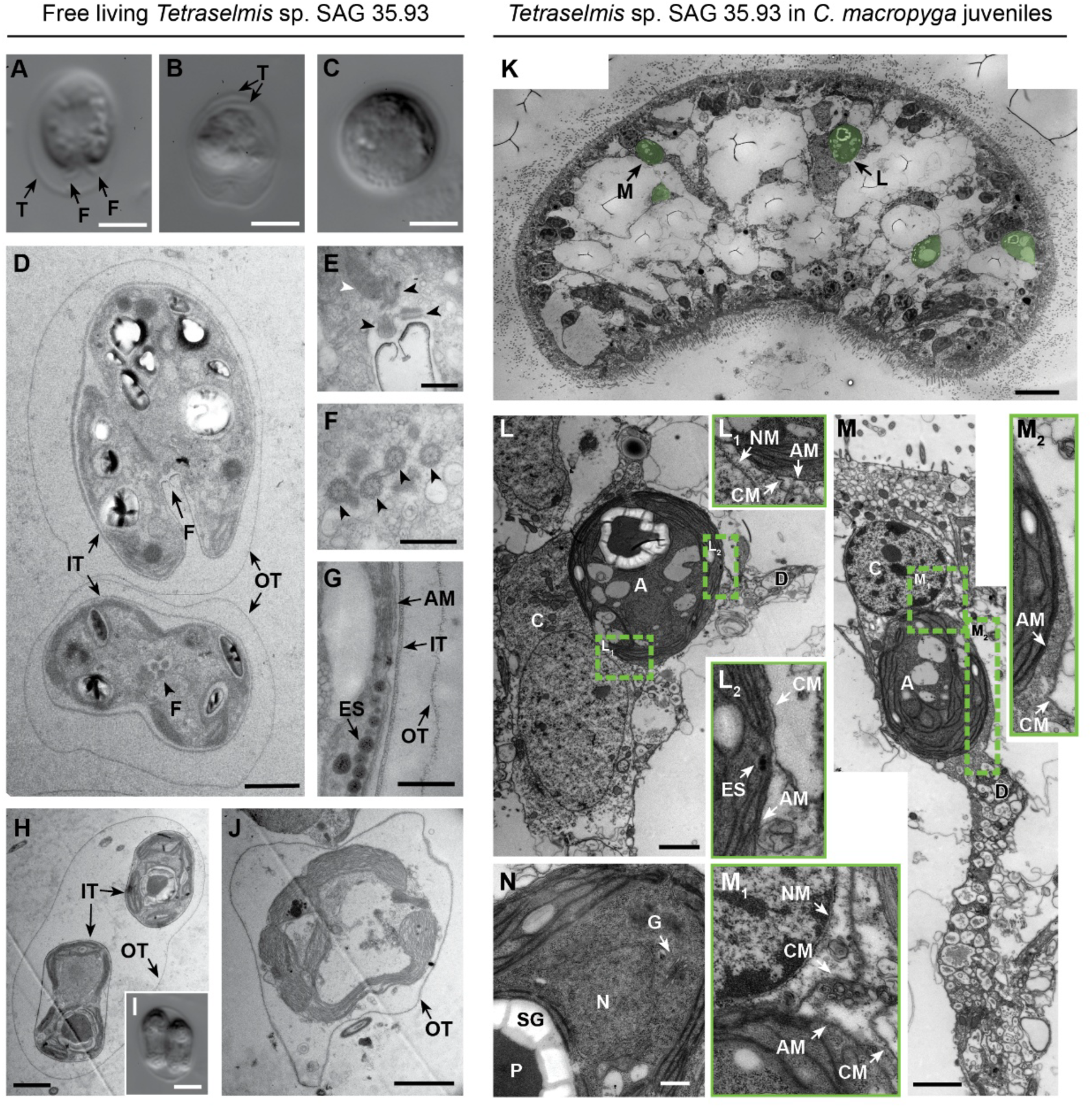
Ultrastructure of *Tetraselmis* algae in culture and within juveniles. *Tetraselmis* sp. SAG 35.93 algae free living from culture (A-J) or inside *C. macropyga* juveniles after uptake (K-N) observed with light microscopy with Differential Interference Contrast (DIC) (A-C,I) or Transmission Electron Microscopy (TEM) (D-H,J-N). **A** Algal cell with two of the 4 flagella visible inside the theca. **B** Algal cell with two layers of theca visible. **C** Algal cell with a young cyst morphology: round and without visible flagella (Kang et al., 2024). **D** Ultrastructure of free-living algae showing the two thecal layers, the flagellar pit with flagellar slit in longitudinal section (F – arrow) and flagella in cross section (F – arrowhead). **E** Flagellar pit with centrioles in the flagellar bases (black arrowheads) and rhizoplast (white arrowhead). **F** Cross section through all 4 flagella (arrowheads). **G** Detail of an algal cell showing eyespot, the plasma membrane, the inner theca, and the outer theca. **H** Daughter cells after cell division still surrounded by the outer theca of the parent algal cell. **I** DIC picture of daughter cells after cell division. **J** Algal cell showing a disrupted morphology with only the outer theca present. **K** *C. macropyga* juvenile cross section in overview, with algae pseudocoloured in green; arrows mark the algae shown in L,M. **L** Algal cell associated with a parenchymal animal cell (C); details of the membranes in **L_1_** and **L_2_**. **M** Algal cell associated with an epidermal animal cell (C) and with a process containing digestive vacuoles (D); details of the membranes in **M_1_** and **M_2_**. **N** Detail of algal cell in parenchyma with visible Golgi apparatus. A: Algal cell, AM: Algal cell membrane, C: Animal cell, CM: Animal cell membrane, D: digestive vacuoles, ES: eyespots, F: Flagellum, G: Golgi apparatus, IT:I Inner Theca, N: nucleus, NM: Nuclear membrane, OT: Outer Theca, P: pyrenoid, SG: starch granules, T: Theca; scale bars are 10 µm in K; 5 µm in **A,B,C,I**; 2 µm in **D,H,I,L,M**; 0.5 µm in **E,F,G,N**.

Upon TEM analysis, we found algal cells in 4 individuals out of 5 *C. macropyga* juveniles exposed 4-5 days to *Tetraselmis* algae. No algae were observed in one control juvenile not exposed to *Tetraselmis*. Confirming our confocal microscopy observations, the algae can associate to the body wall, but they are mostly found in the parenchyma; they can be intact and round-shaped, but they can also show signs of damage (Fig. 3L,M, Fig. S3A,B,C). Although algae can be adjacent to epidermal cells, they are enclosed by a membrane and surrounded by digestive vacuoles, likely belonging to a different animal cell (Fig. 3L,L_2_,M,M_2_). All the observed algae inside *C. macropyga* juveniles have neither a theca nor flagella (Fig. 3K,L,M), though flagellar bodies are found in one instance (Fig. S3D). Eyespots and Golgi apparatus are visible (Fig. 3L_2_,N).

### In adults, symbionts are extracellular at the body wall, but can be intracellular in the parenchyma

Adult histological sections show that algal symbionts are either associated with the body wall—under the muscle layer or between the epidermis and the muscle layer—or alternatively, they are located inside the body, in the parenchyma (Fig. 4). Symbiont density is higher in the body wall than in the parenchyma (Fig. 4B) and, within the body wall, higher in the dorsal than in the ventral side (Fig. 4C). The symbionts are present in any region apart from the digestive system, although they are occasionally at its periphery. They can be closely associated with both male and female gonads, in direct contact with sperm cells and oocytes, as well as adjacent to the nervous system, both the anterior neuropil and the longitudinal nerve cords (Fig. S4).

**Fig. 4.**
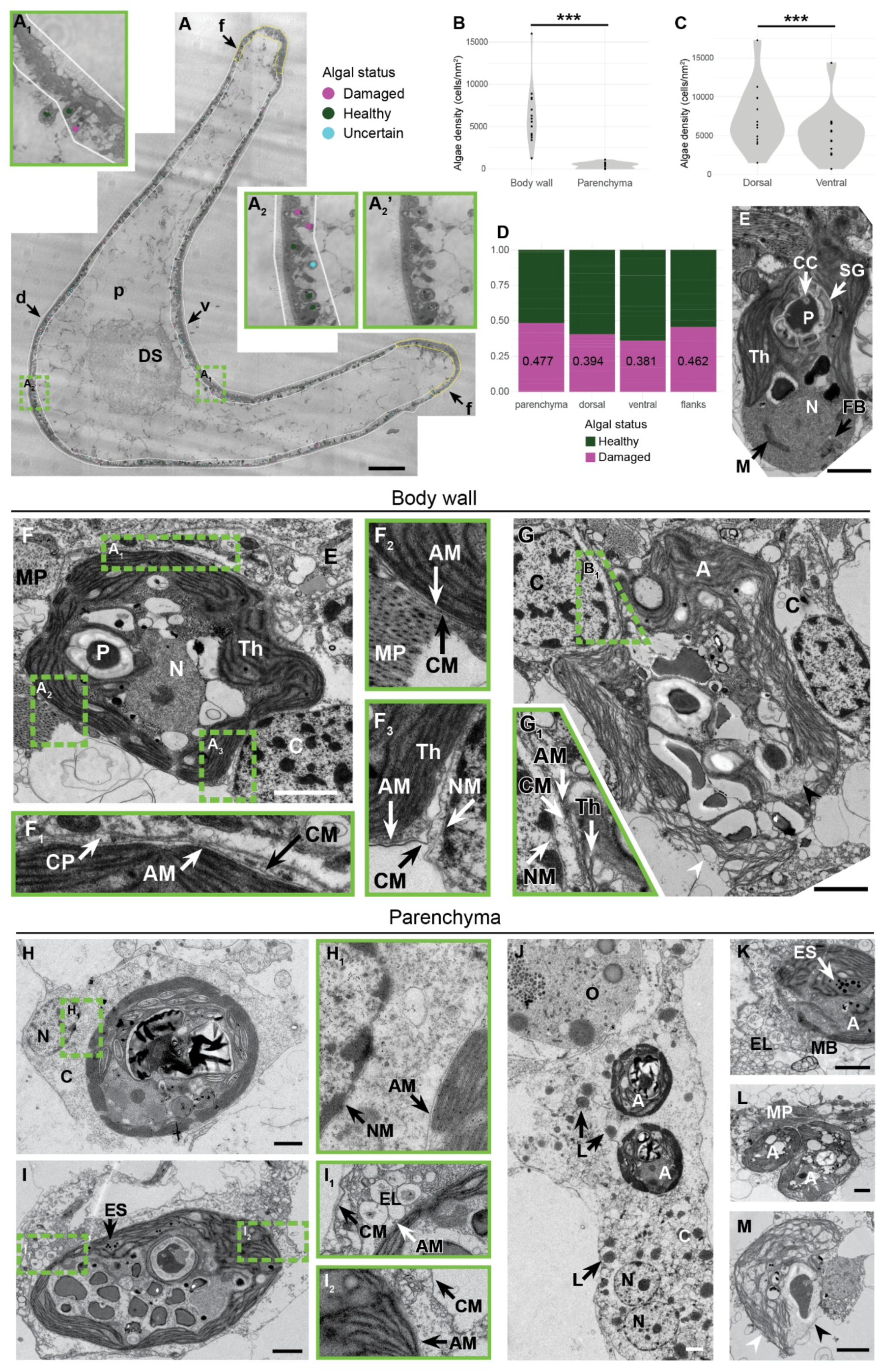
Distribution and cellular interaction of algal symbionts within *C. macropyga*. **A** Cross section imaged with coherence contrast, the regions considered for the analysis are bordered in white or yellow and marked as p: parenchyma, v: body wall – ventral, d: body wall – dorsal, f: body wall – flanks (in yellow); algae are marked by coloured dots as healthy, damaged, and uncertain (green, magenta, and cyan, respectively). **A_1_, A_2_, A_2_’** higher magnification of the dashed rectangles in **A**. **A_2_’** shows the same area as **A_2_** without annotations. **B** Symbiont density in body wall and parenchyma (Wilcoxon signed rank exact test, V = 0, p-value = 0.0001221, n = 14; mean ± SE (cells/nm^2^) for body wall: 6286.62 ± 943.44; for parenchyma: 413.38 ± 93.24). **C** Symbiont density in dorsal and ventral sides of the body wall (Wilcoxon signed rank exact test *wilcox.test*: V = 104, p-value = 0.0002441, n = 14; mean ± SE (cells/nm^2^) for dorsal side: 7260.32 ± 1051.31; for ventral side: 5221.81 ± 872.53). **D** proportion of damaged and healthy symbionts in the different regions (generalised linear model *glmer*, with binomial distribution function; maximal model: cbind(Num.Damaged_algae,Num.Healthy_algae) ∼ region + (1|sample/section); minimal adequate model: cbind(Num.Damaged_algae, Num.Healthy_algae) ∼ 1 + (1 | sample/section); n=14). The median of the proportion of damaged algae for each region is shown on the column. **E-M** TEM photographs of symbiotic algae: **E** shows the ultrastructural morphology. **F, G** symbionts associated to the body wall, between the epidermis and the muscle layer or under the muscle layer (oriented toward the top of the image); **H-M** symbionts in the parenchyma. In **F_1_, F_2_, F_3_, G_1_, H_1_, I_1_, I_2_** details of the membranes showing the intra- or extracellular status of the symbionts. **G, M** show damaged symbionts; white arrowhead points at thylakoids with a disrupted structure, black arrowhead at a collapsing pyrenoid. A: Algal cell, AM: Algal cell membrane, C: Animal cell, CC: cytoplasmic canaliculi, CM: Animal cell membrane, CP: Cell processes, DS: digestive system, E: epidermis, EL: Endolysosome, ES: Eyespot granules, FB: Flagellar Bases, L: Lipid bodies, M: Mitochondrion, MB: Multilamellar body, MP: Muscle cell processes, N: Nucleus, NM: Nuclear membrane, O: oocyte, P: pyrenoid, SG: Starch granules, Th: Thylakoids; *** : p<0.001; Scale bars are 100 µm in **A**, otherwise 2 µm.

Some symbionts exhibited morphological degradation, including loss of thylakoid structural integrity, nuclear disintegration, pyrenoid collapse, and plasma membrane disruption (Fig. 4G,M). Based on these features, symbionts were categorized as healthy, damaged, or uncertain (Fig. 4A,A2,A2’). If symbionts were retained in the body wall for photosynthetic function and internalized for digestion, healthy algae would be expected to predominate in the body wall and damaged algae in the parenchyma, as reported for *Convoluta convoluta* (Pedersen, 1964). However, no significant regional differences in the ratio of damaged to healthy symbionts were detected (Fig. 4D).

The association between algal symbionts and host cells was examined using TEM in ∼200 algae from three adult individuals. The matrix of the pyrenoid, a carbon-fixation organelle (He et al., 2023), was surrounded by starch grains and penetrated by cytoplasmic canaliculi (Fig. 4E). The nucleus was round and close to mitochondria (Fig. 4E). Eyespots were present, whereas thecae, flagella, and the flagellar pit were absent; flagellar bases were observed in two instances (Fig. 4E,I,K).

The algae associated to the body wall are always extracellular (Fig. 4F_1_-F_3_,G_1_). In the parenchyma, we find symbionts extracellularly (Fig. 4L,M), but they can also be intracellular (Fig. 4H-K). While the shape of the extracellular symbionts is irregular, as in the body wall, intracellular algae are always rounder (cf. Fig. 4F,G,L and Fig. 4H,I,J,K). Even though no symbionts are found in the digestive system, algae-bearing cells show characteristics typical of acoel digestive cells, i.e. they are vacuolated (Fig. 4I,K) and multinucleated (Fig. 4J), and they contain endolysosomes, multivesicular bodies, and lipid bodies (Fig. 4I,J,K) (Gavilán et al., 2019; Hartenstein & Martinez, 2019; Jastrow, 2021; J. P. S. Smith, 1981). These algae-bearing cells can also be in direct contact with oocytes (Fig. 4J, Fig. S4).

### Pathways and processes enriched in photosymbiotic adults versus aposymbiotic juveniles

To identify changes in post-embryonic development and photosymbiosis maintenance at a molecular level, we performed bulk transcriptomics on *C. macropyga* aposymbiotic juveniles and symbiotic adults. The samples are grouped by ontogenetic stage in hierarchical clustering (Fig. S5A), whereas in principal component analysis the juvenile samples all cluster very closely together and the adult samples present more variability (Fig. S5B). More genes are upregulated than downregulated in adults: at adjusted p value < 0.001, the 3.3% (21 944 out of 66 1165 genes with non-zero read count) is upregulated (log2 Fold Change > 1), and the 1.1% (7 472 out of 66 1165) is downregulated (log2 Fold Change < -1) (Fig5A). We used Gene Ontology (GO) and Kyoto Encyclopedia for Genes and Genomes (KEGG) annotations to perform Gene Set Enrichment Analyses (GSEA): out of 699 959 genes in *C. macropyga de novo* assembled transcriptome, 32 845 (4.7%) are annotated for GO terms and 13 961(2.0%) for KEGG modules.

**Fig. 5.**
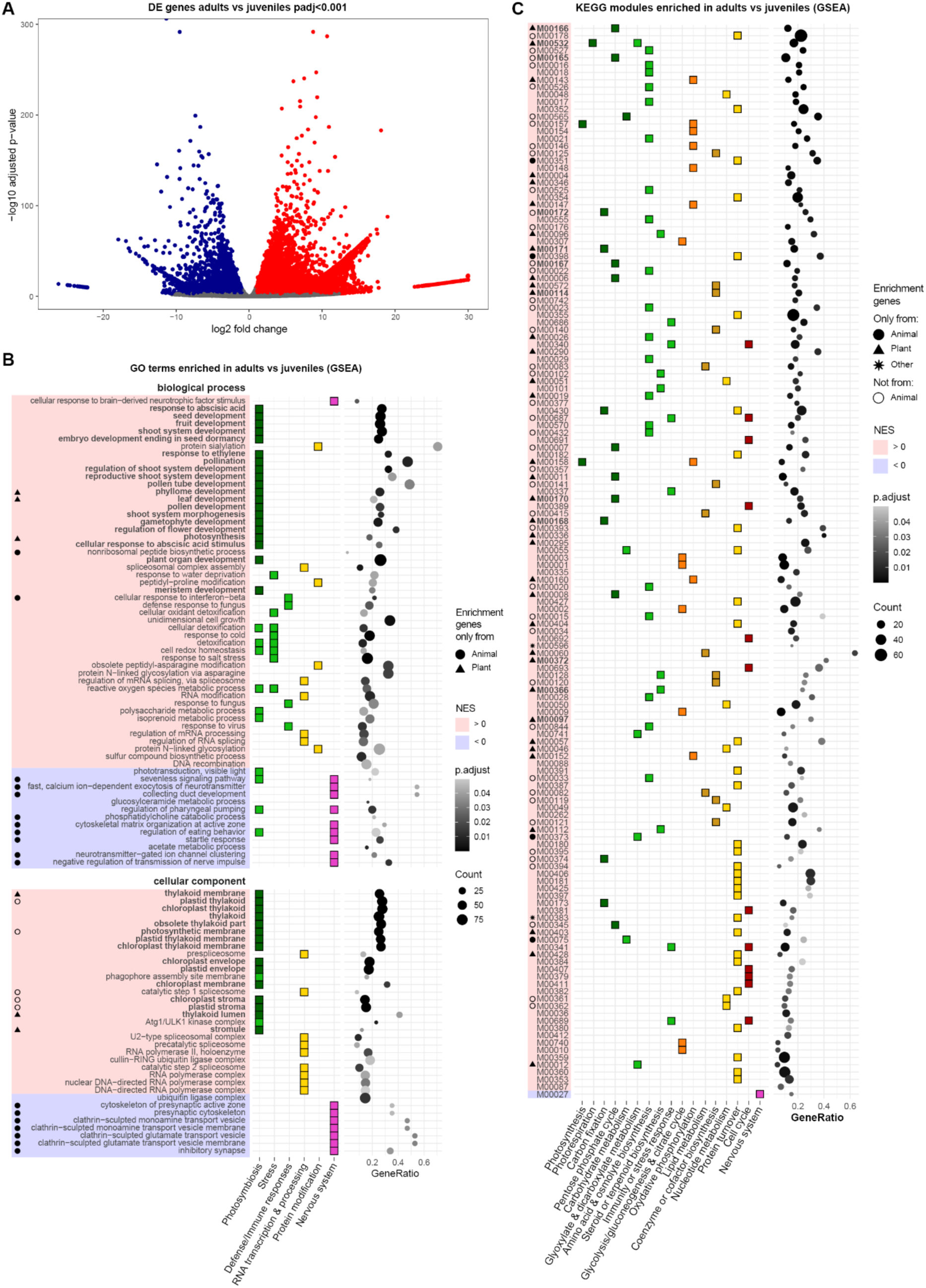
Transcriptomic analysis. Differential gene expression in *C. macropyga* adults compared to juveniles. **A** Volcano plot of differentially expressed genes. Blue dots represent downregulated genes in adults and red dots upregulated genes (padj <0.001, absolute value of log2 fold change > 1). **B,C** Gene Set Enrichment Analysis of genes ranked by log2fC*(-log10 p) with GO terms (**B**) or KEGG modules (**C**). Gene sets are ordered by Normalized Enrichment Score (NES), with positive enrichment (activation) in red and negative enrichment (suppression) in blue. Dark green: plant gene sets, light green: gene sets potentially involved in photosymbiosis. Gene sets in bold are specific for photosynthetic organisms. Names corresponding to KEGG modules in (**C**) can be found in Table S3.

GSEA reveal a positive enrichment for various plant-related processes and cellular components (Fig. 5BC, Table S1-S4). For most of the positively enriched gene sets, core enrichment genes belong not only to Metazoa or Viridiplantae, but also to other groups, such as Fungi, SAR, Bacteria, Archaea (Fig. 5BC, Table S1-S4). This could mean that various symbiotic partners all regulate their genes individually or that they all contribute to a single process in the holobiont; however, it is possible that the genes belong to *C. macropyga* or *Tetraselmis* sp., while the blastx search used to annotate them retrieves genes from a different clade.

Negatively enriched gene sets are mostly related to the nervous system, and almost all core enrichment genes contributing to them belong to Metazoa (Fig. 5BC, Table 1-S4). Among the positively enriched we find processes and pathways associated with photosynthetic organisms: photosynthesis, plant organ development, responses to plant hormones (GO biological processes, Fig. 5B); photosynthesis, carbon fixation, photorespiration, the pentose phosphate cycle, beta-carotene biosynthesis (KEGG modules, Fig. 5C). Other positively enriched categories are potentially associated with photosymbiosis: responses to stress, especially oxidation; immune responses; phagophore membranes; biosynthesis of steroids, terpenoids, amino acids or osmolytes; glyoxylate and dicarboxylate metabolism; carbohydrate and lipid metabolism. Other categories of positively enriched sets could be either related to photosymbiosis or development and differences between ontogenetic stages: energy-producing pathways (glycolysis, citrate cycle, oxidative phosphorylation); protein turnover (RNA transcription, processing, translation, and degradation, ribosomes, protein processing, modification, and degradation); metabolism of nucleotides; coenzymes and cofactor biosynthesis; cell cycle regulation.

## Discussion

Our data provides a characterisation of the association between *Convolutriloba macropyga* and its endosymbiotic green algae from a morphological, molecular and developmental perspective. Moreover, from Rubisco phylogeny, the symbionts are identified as belonging to *Tetraselmis* and closely related to *Convolutriloba longifissura*’s symbionts (Fig.1 CD).

### Acoel development and horizontal symbiont acquisition

Embryonic development is faster in *Convolutriloba macropyga* than *Hofstenia miamia, Symsagittifera roscoffensis*, or *Neochildia fusca* (Arboleda et al., 2018; Henry et al., 2000; Kimura et al., 2021). This could be linked to them being kept at room temperature in the laboratory, while *C. macropyga* is cultured at warmer, tropical-like, temperatures. *C. macropyga* follows the acoel stereotypical duet cleavage (Hejnol, 2015) and starts gastrulation at 14-cell stage, slightly earlier than *H. miamia* and *S. roscoffensis* (16-cell stage (Bresslau, 1909; Kimura et al., 2021)). When, at 3 days post egg laying, *C. macropyga* juveniles hatch, they are drop-shaped and devoid of algal symbionts (Fig. 1, (Shannon & Achatz, 2007)). They swim actively and, when provided with *Tetraselmis* algae, ingest them through the mouth (Fig. 1, Video S1,2,3). When *C. macropyga* juveniles are exposed to *Tetraselmis* prospective symbionts, they start taking them up straight away (Fig. 1F, 2BC). Slightly more than a quarter of the juveniles shows no algal symbiont, regardless of the exposure time (Fig. 1F); by comparison, the ratio of *S. roscoffensis* taking prospective symbionts up depends on the concentration of algae in the medium (Douglas, 1983). The number of algae within juvenile increases over time both in *C. macropyga* and *S. roscoffensis*, even after *Tetraselmis* algae are removed from the culture medium (Fig. 2BC, (Douglas, 1983)). This strongly supports proliferation of the algae within acoel juveniles.

As soon as *C. macropyga* juveniles start taking the algae up, these can be found associated to the body wall (Fig. 2EF). Conversely, in *S. roscoffensis,* symbionts at the worm periphery are only observed at 11-12 days after ingestion, and not at 2-3 days (Douglas, 1983). In terms of algal spatial dynamics, we do not detect any significant changes in position on the left-right and anterior-posterior axes (Fig. S1BC). However, the algae are displaced along the dorso-ventral axis (Fig. 2D, S1D), with a stark preference for the dorsal side (Fig. S1E), consistently with their position in adults (Fig. 4C). Douglas (1983) reports both athecate and thecate algae in *S. roscoffensis* juveniles upon symbiosis establishment. Although our methods do not allow a thorough analysis, all the observed algae in *C. macropyga* juveniles are athecate (Fig. 3K,L,M, Fig. S3).

Together with the theca, *Tetraselmis* lose flagella—but not flagellar bodies—upon uptake (Fig. 3, Fig. 4), similarly to *S. roscoffensis* and *C. longifissura* (Hirose & Hirose, 2007; Oschman, 1966). However, eyespots are retained in *C. macropyga* symbionts (Fig. 3E,G, Fig. 4).

### Amino acid synthesis, oxidation of fatty acids, stress responses, and osmoregulation are enriched in photosymbiotic adults

Once algae and hosts recognize each other and the symbiosis is established, specific mechanisms are needed to maintain it (Davy et al., 2012). Our transcriptomic analysis comparing photosymbiotic adults and aposymbiotic juveniles aims at elucidating processes relevant for symbiosis maintenance (Fig. 5). While some enriched gene sets most likely reflect the difference in ontogenetic stage, other adult-specific pathways or processes could play a role in photosymbiosis (Fig. 6).

**Fig. 6.**
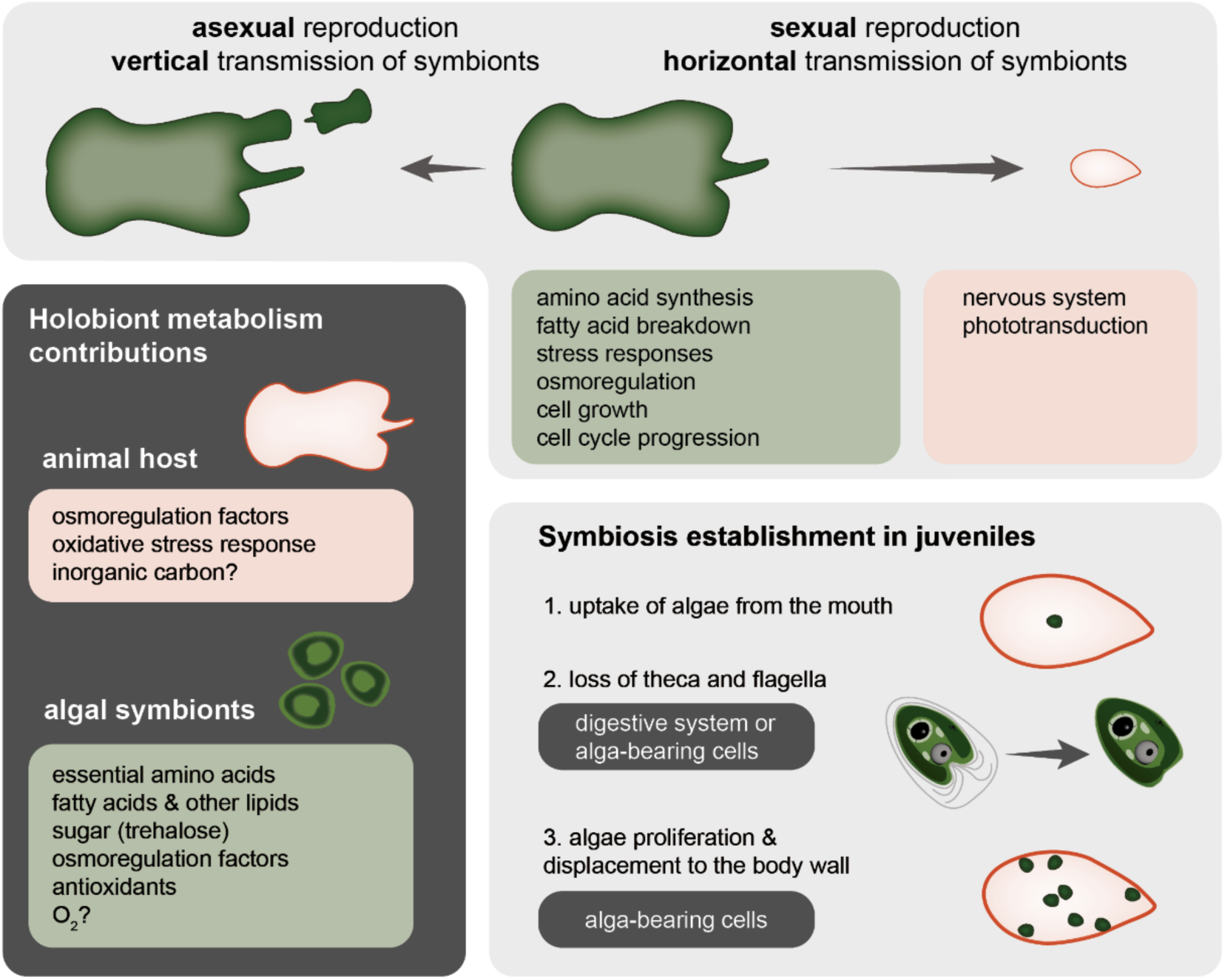
Metabolic interactions and symbiosis establishment. Transmission modes of *Tetraselmis* sp. symbionts in *C. macropyga*, metabolic interactions between the symbionts and the host, and establishment of symbiosis in juveniles. Interpretations and working model on dark grey background.

Many enriched processes relate to protein modification (sialylation, glycosylation, amino acid-specific modification): some could be important for interaction with symbionts (Li & Ding, 2019). Phagophore membranes are an enriched GO cellular component, suggesting that autophagy might play a role in symbiosis or other adult-specific processes.

Beta oxidation of fatty acids is enriched in adults, as well as the glyoxylate pathway and the ethylmalonyl cycle. Moreover, the KEGG modules for fatty acid biosynthesis (M00415, M00083, M00083) are enriched with almost no core enrichment genes annotated to metazoans (Table S3,S4). This suggests that photosymbiotic adults break down fatty acids and use them as a source of energy (Alber, 2011; Nelson et al., 2013), as proposed for photosymbiotic sea anemones receiving fatty acids from dinoflagellates (Lehnert et al., 2014; Matthews et al., 2018). Sterols are also potentially transferred by photosymbionts (Fig.5, Table S3,S4) (Yee et al., 2025). Carbohydrate metabolism modules are also enriched, with genes from plants for the biosynthesis of trehalose, and from plants and animals for the production of N-glycans (for protein modification). In cnidarians, photosymbionts transfer fixed carbon as glycerol, glucose, fumarate, succinate, or maltose (Davy et al., 2012). In acoels, trehalose or its derivatives could be transferred instead.

In the coral-dinoflagellate symbiosis, dinoflagellates are responsible for most of the nitrogen fixation (Pernice et al., 2012) and for the synthesis and translocation to the host of essential amino acids (Lehnert et al., 2014; Wang & Douglas, 1999). Since many KEGG modules involved in the biosynthesis of amino acids are activated in photosymbiotic *C. macropyga* adults, the photosymbionts may be providing amino acids to the host. In particular, many of the pathways activated concern essential amino acids not produced by animals, with the exception of proline and cysteine (Table S3,S4, (Kasalo et al., 2024)). Essential amino acid translocation by the symbionts happens also in *S. roscoffensis* (Boyle & Smith, 1975).

The amino acids synthesised from enriched pathways might also contribute to protection against UV (Muller-Parker et al., 2015) or to osmotic homeostasis (Mayfield & Gates, 2007). In fact, free amino acids are candidate osmoregulation factors in cnidaria-dinoflagellate symbiosis—although cnidarian endosymbionts are strictly intracellular and release the amino acids into the cell cytoplasm (Mayfield & Gates, 2007). Moreover, betaine is a known osmolyte in marine environments and photosymbiosis (Diaz et al., 1992; Yancey et al., 2010). Responses to water deprivation and to salt stress are also enriched processes in the adults (Fig. 5, Table S1). Together, these gene sets suggest an increased effort towards osmoregulation in adults.

Endosymbiont photosynthesis brings about the problem of accumulation of O_2_ and Radical Oxygen Species (ROS) in animal tissues (Ip & Chew, 2021). It is therefore not surprising the enrichment of processes dealing with ROS metabolism, antioxidant activity, cellular oxidant detoxification, and cell redox homeostasis. Furthermore, *Tetraselmis* species themselves synthesise antioxidants, such as tocopherols (Conlon & Touzet, 2025), a process enriched in photosymbiotic *C. macropyga* adults, too. Moreover, a few GO terms and KEGG modules pertaining to immunity are enriched: they are mostly related to stress and inflammation or to antiviral immunity, which are linked to oxidative stress (Fig. 5, Tables S1, S3) (Shang & Taylor, 2011; Zhang et al., 2024).

Many processes activated in *C. macropyga* adults are linked to growth, from RNA and protein synthesis and modification to cell cycle transitions (Fig. 5, Tables S1, S3). A likely explanation is the growth and asexual reproduction of adults, which obtain nutrients both from the endosymbionts and from feeding. Juveniles were not fed since we have not identified any food they are able to ingest yet. Few of the core enrichment genes related to growth and cell cycle are plant genes, which suggests the algal symbionts undergo cell division. However, electron microscopy does not show algae dividing. The absence of symbiont cell division in adults, but not in juveniles, is also reported for *S. roscoffensis* (Arboleda et al., 2018), although it could be restricted to the night-time (Oschman, 1966). The enrichment of processes from the citrate cycle and glycolysis is likely due to an increase in oxygen and nutrient availability; it results in the increase of energy available (Arnold & Finley, 2023).

### Intracellular symbionts in parenchymal cells: digested or transported?

Keeping the symbionts within organs or in symbiosomes inside cells is usually important to prevent interference with host processes and it can create a favourable environment for the endosymbionts (Chomicki et al., 2020; Fronk & Sachs, 2022; Liao et al., 2025). E.g., in intracellular cnidarian-dinoflagellate photosymbiosis, the symbiosome is acidified to promote photosynthesis (Chomicki et al., 2020; Liao et al., 2025), in giant clams the tubular system harbouring extracellular photosymbionts is also kept optimal for photosynthesis (Ip & Chew, 2021).

Algal symbionts in *C. macropyga* are not restricted to specific organs or tissues. In fact, they are mostly located at the body wall, but also in internal tissues, even close to nervous and reproductive systems (Fig. 4, Fig. S4). Moreover, the majority of them are extracellular, free of membrane-bound symbiosomes (Fig.4). This suggests no need for physical containment, which can be explained in two ways. (1) The host has non-physical mechanisms of controlling the algal cell cycle and proliferation; this seems supported by the unchanged algae to animal cell ratio in the presence of potential symbiotic algae and prey (Fig. S2). (2) the host is farming the endosymbionts and therefore needs to maximise their number and distribution. This interpretation is coherent with the fact that the symbiosis is obligate for the animals, but facultative for the algae, i.e. algae can be isolated and found free living, but adult animals are never aposymbiotic and die when kept in the dark (Shannon & Achatz, 2007). Besides, the photosynthetic rate is lower in *Tetraselmis convolutae* within *S. roscoffensis* than in free-living ones (Androuin et al., 2020).

Whereas photosymbionts associated to the body wall are always extracellular, they are sometimes observed within cells in the inner parenchyma (Fig. 4). Extracellular symbionts mainly have a flexible shape, adjusted to the neighbouring cells, but intracellular symbionts are rounder (Fig. 4). Although this duality has not been described in *S. roscoffensis*, it remains possible—if not likely: intracellular algae are mentioned along with extracellular ones by (Oschman, 1966); Fig. 3 in (Bailly et al., 2014) shows both irregularly-shaped symbionts close to the body wall and rounder, potentially intracellular, symbionts near oocytes and sperm.

The irregular shape of green algal symbionts in acoels has been attributed to the loss of a theca (Liao et al., 2025), yet the absence of a theca is compatible with a round shape of the algae too (Fig. 4, and (Hikosaka-Katayama et al., 2024)). The adjustment of symbiont morphology to the neighbouring cells it could denote their close interaction (Hikosaka-Katayama et al., 2024), for example in exchanges with host cells (Fig.7). It could also be due to physical properties of host cells or extracellular matrix.

**Fig. 7.**
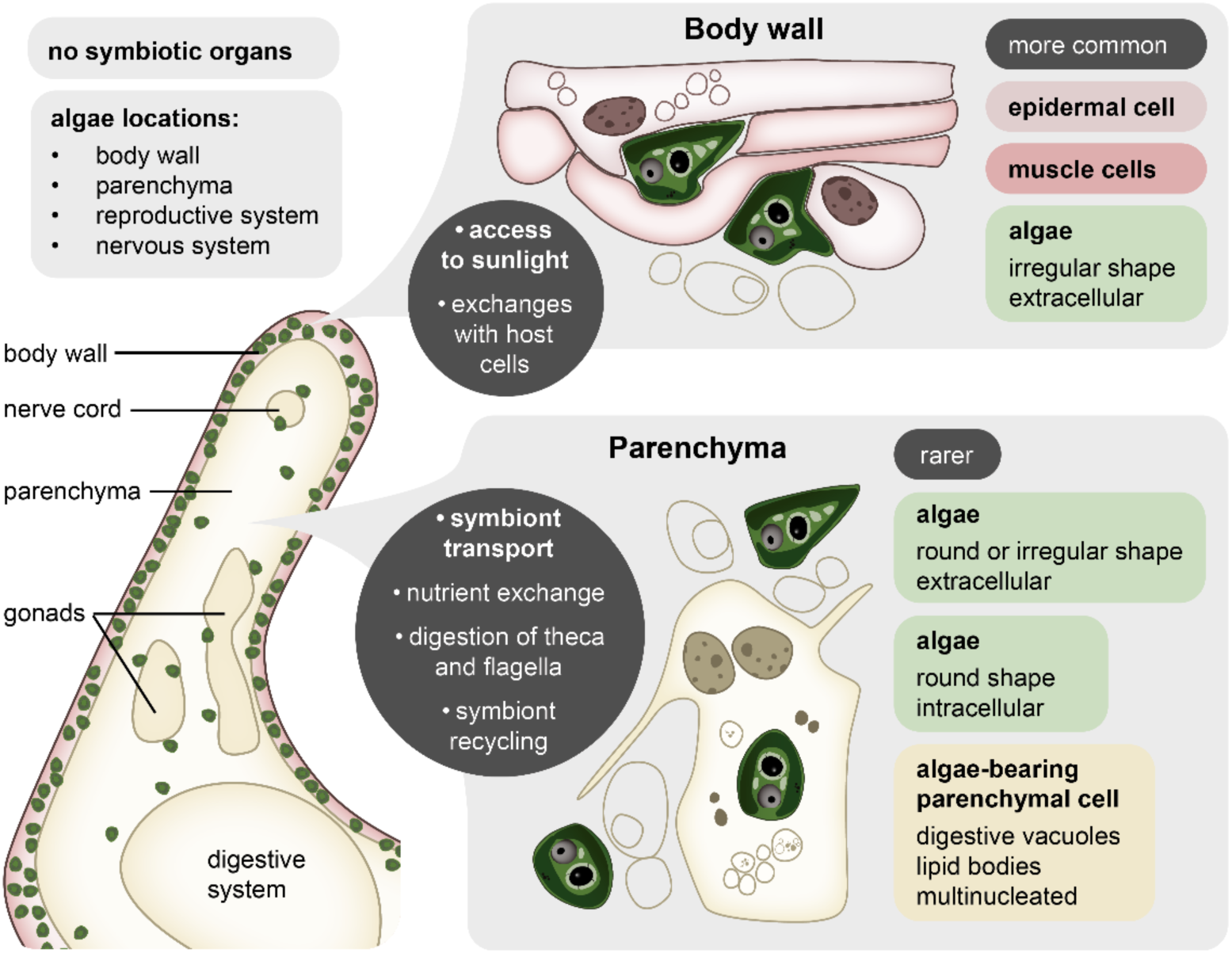
Symbiont location and interaction with host cells. Schematic view of a cross section of *C. macropyga* with location of the algal symbionts and their interaction with host cells. Working model on dark grey background, with in bold the interpretation we deem most likely.

Since the majority of symbionts is extracellular and associated to the body wall, we regard this as the most stable state of *C. macropyga-Tetraselmis* sp. symbiosis. They are mostly at the dorsal body wall, which is more easily accessible to sunlight. The less abundant symbionts intracellular in the parenchyma could be interpreted as (1) host cells acquiring nutrients, (2) newly ingested algae that are being stripped of their theca and flagella, (3) symbionts being transported (Fig.7). These options are discussed in detail below.

1. The presence of digestive vacuoles and other digestive cell features in the cells with intracellular symbionts (Fig. 4) suggests they could be obtaining nutrients from them, either by digesting photosynthates or by digesting the symbionts themselves, as in other types of symbiosis (Sogin et al., 2020; Yee et al., 2025). Notably, the presence of lipid bodies is consistent with the transfer of fatty acids suggested by transcriptomic analyses. However, damaged symbionts are equally present in the parenchyma and in the body wall (Fig. 1D) and many of the algae within parenchymal cells are intact (Fig. 4H,I,J,K), which do not favour the hypothesis of intracellular digestion of the symbionts in parenchymal cells.
2. The loss of algal theca and flagella in juveniles most likely happens inside the algae-bearing cells or in the digestive system. Nonetheless, this cannot be the only function of these host cells since our data do not support a continuous uptake of symbionts by adults (Fig.S2). Besides, the animals were isolated two days before fixation for TEM and had no access to potential symbionts from the environment. It is therefore unlikely that the intracellular algae in the adult parenchyma are newly acquired symbionts being processed.
3. In the acoel *Waminoa sp.,* intracellular dinoflagellate symbionts are extremely mobile, thanks to host cell processes (Zhong et al., 2025). Chlorophyte symbionts could be transported to different body regions in *Convolutriloba* too, with the difference that this requires a transition between intra- and extracellularity. This could be important to exchange metabolites or find optimal light conditions (Yee et al., 2025). In this scenario, we can interpret the algae within parenchymal cells close to the body wall in juveniles (Fig. 3M) as in the process of being transported to the periphery of the body for symbiosis establishment.

Based on the presence of digestive vacuoles, the algae-bearing parenchymal cells could be phagocytic cells, not belonging to the digestive system. Regardless of the process, the animal cells containing the symbionts likely belong to the peripheral parenchyma described in acoels: cells surrounding the digestive system (central parenchyma) (Gavilán et al., 2019; J. P. S. Smith, 1981). Among the proposed functions for peripheral parenchymal cells in acoels are phagocytosis and the distribution of nutrients (Gavilán et al., 2019), which fit well their co-option to digest photosynthates, theca and flagella, or whole symbionts, or to transport them throughout the host.

## Conclusions

In this work, we study the cellular interactions between the acoel *Convolutriloba macropyga* and its photosymbionts, the chlorophyte algae *Tetraselmis* sp, which can be transmitted both vertically and horizontally. We describe the symbiont uptake and their dynamics within juveniles, as well as their location in juveniles and adults. Our data suggests algae uptake occurs continuously in juveniles but ceases at a later stage in adults, that newly acquired symbionts in juveniles proliferate and are transported to the body wall. In adults, algal symbionts can be extracellular at the body wall and intracellular in algae-bearing parenchymal cells, which might be responsible for their digestion or transportation. Transcriptomic data analyses point at various processes potentially involved in photosymbiosis maintenance, such as translocation of lipids, sugar, and amino acids from the endosymbionts to the host, and responses to osmotic and oxidative stresses. Our findings contribute to the understanding of host-microbe interactions in photosymbiotic associations. They provide a starting point for comparative and evolutionary research aimed at understanding the widespread phenomenon of animal photosymbiosis. Moreover, research on extracellular photosymbionts spread throughout animal host tissues, like in *C. macropyga,* might yield important results for transplants and allorecognition, as well as targeted oncotherapy or vascular perfusion using microalgae (Ehrenfeld et al., 2023; Liu et al., 2024; Vargas-Torres et al., 2024).

## Materials & Methods

### Animals and algae

*Convolutriloba macropyga* Shannon & Achatz 2007 (Shannon & Achatz, 2007) were kept at 26°C with a cycle of 10-14 hours darkness-light in artificial sea water with salinity 36‰ (Classic Sea Salt by Tropic Marin, Dr. Biener GmbH, Wartenberg, Germany); they were fed twice a week with nauplii of brine shrimp *Artemia* sp. Juveniles were collected by isolating freshly laid egg clusters in order to keep them aposymbiotic until fixation or exposure to *Tetraselmis* algae.

Isolation and culture of symbionts from adult *C. macropyga* could not be replicated with methods commonly used in acoels (Boyle & Smith, 1975; Shannon & Achatz, 2007). Therefore, exposure experiments were carried out with *Tetraselmis* sp. algae isolated from *Convolutriloba longifissura* (Bartolomaeus & Balzer, 1997) and commercially available (Culture Collection of Göttingen University, SAG strain 35.93) (Balzer, 1999). They were kept at room temperature in sea water medium (SWES = “Seewasser + Erddekokt + Salze”) prepared according to SAG instructions (Culture Collection of Algae (SAG), n.d.) and subcultured every 4-8 weeks.

### Phylogeny of symbionts and free-living *Tetraselmis* algae

RNA was extracted with Trizol and 1–bromo–3–chloropropane from *C. macropyga* adults and free-living *Tetraselmis* sp. SAG 35.93 and cDNA synthesis was performed with ABScript II cDNA First-Strand Synthesis Kit (ABClonal, Woburn, MA, USA), following the manufacturer instructions and using random primers. Partial chloroplast rbcL genes were amplified with PCR (primers 5′-TAGGTYTWTCAGCTAAAAACTACGG-3′ and 5′-GCTGGCATGTGCCATACGTGAATAC-3′ (Hikosaka-Katayama et al., 2024)); nuclease-free water was used as negative control. The PCR conditions were slightly modified from (Hikosaka-Katayama et al., 2024): initial denaturation (94°C, 60 s), 35 cycles (denaturation – 98°C, 10 s, annealing – 52°C,5 s, and extension – 68°C, 5 s), final extension (68°C, 5 min). PCR products were checked by gel electrophoresis on 1% agarose gel in TAE (40 mM Tris, 20 mM acetic acid, 1 mM EDTA) and Sanger sequencing was carried out by Eurofins Genomics Europe Shared Services GmbH (Ebersberg, Germany). The sequences were aligned with other rbcL sequences from algal and cyanobacteria using MAFFT v7.526 with L-INS-I strategy (Katoh & Standley, 2013), trimmed with TrimAl v1.5.rev0(Capella-Gutiérrez et al., 2009); phylogenetic analysis was performed with iqtree2 v.2.4.0 (Minh et al., 2020), with substitution model GTR+F+G4 chosen with ModelFinder (Kalyaanamoorthy et al., 2017) and node support evaluated from 1000 iterations of ultrafast bootstrap (Hoang et al., 2018).

### Exposure of juveniles and adults to *Tetraselmis* algae

Egg clusters were collected from the aquarium and transferred to Petri dishes containing clean filtered artificial seawater (FASW). The isolated eggs were inspected daily, and hatchlings were transferred to a new Petri dish. Juveniles were maintained in algae-free FASW for up to five days post-hatching. Subsequently, exposure to the algal strain *Tetraselmis* SAG 35.93 was initiated by adding 1 ml of algal suspension prepared from culture tubes. Prior to exposure, the algal suspension was centrifuged, and the pellet was resuspended in fresh FASW. Following the exposure period, juveniles were once again isolated from the algae until fixation (Post Exposure Days, PED). When fixation occurred on the same day as exposure, it was carried out immediately after algae removal. For subsequent fixation time points, samples were processed at approximately 24-hour intervals to ensure comparability across PED conditions.

*C. macropyga* adults were cleaned twice with filtered artificial sea water, placed in a well of a 12 well plate in 2 ml of filtered artificial sea water (fASW). 60 individuals were used as control, *Tetraselmis* algae SAG 35.93 were added to 60 individuals, and another 60 individuals were fed with *Artemia* sp. nauplii every 5 days. The day after feeding, the medium was changed for samples in all conditions. After 19 days, the animals were fixed for imaging.

### Light microscopy of developing *C. macropyga* and *Tetraselmis* algae

Developing embryos were mounted on a slide and covered with a coverslip on clay feet. Images were taken with Zeiss Axiocam HRc connected to a Zeiss Axioscope Ax10 (Carl Zeiss Microscopy GmbH, Jena, Germany) using bright-field Nomarski optics. Living juveniles exposed to algae were imaged and filmed through a Leica DM750 equipped with a Leica ICC50 W camera (Leica Microsystems, Wetzlar, Germany). Free-living SAG 35.93 algae were mounted on a slide with a coverslip on clay feet. They were imaged at 400x magnification through a customised Axio Imager.M2 (Carl Zeiss Microscopy GmbH, Jena, Germany) equipped with a pco.panda sCOMS camera (Excelitas PCO GmbH, Kelheim, Germany). Colour videos were taken with an Axiocam 503 color on an Axioscope 5 with DIC (Carl Zeiss Microscopy GmbH, Jena, Germany).

### Confocal Laser Scanning Microscopy of algae in exposed juveniles and adults

*C. macropyga* were washed with fASW, relaxed in 1:1 7% MgCl_2_:fASW (adults) or by adding several drops of 7.5% MgCl₂ (juveniles), fixed with 16% paraformaldehyde diluted 1:4 in fASW for 30 min at room temperature, washed 5x in PBS (Phosphate Buffer Saline). Adults were stained for 30 min with 2.5 µg/ml Hoechst 33342 at room temperature in the dark, washed 2x in PBS, and mounted in 70% glycerol in PBS. Juveniles were incubated in 0.66 µM phalloidin in PBS at room temperature, for 20 or 15 minutes, when using Alexa Fluor™ 488 Phalloidin or Alexa Fluor™ 555 Phalloidin (Invitrogen, Thermo Fisher Scientific, Waltham, MA, USA), respectively. They were mounted in Fluoroshield mounting medium containing DAPI (Sigma-Aldrich, Merck KGaA, Darmstadt, Germany). A LSM 980 confocal laser scanning microscope equipped with an Axiocam 305 mono ZEISS camera (Carl Zeiss Microscopy GmbH, Jena, Germany) was used to image Chlorophyll B autofluorescence and staining signal in whole juveniles and in a flat area between the eyespots and the mouth of adults.

Algae were manually counted and scored in Fiji (v2.16.0) for juveniles (Schindelin et al., 2012). The dorsal or ventral positioning was determined using the focal plane corresponding to the mouth as centre. Algae on the same focal plane as the mouth were not included in any of the two groups. Algae associated with the body wall were quantified by delineating the wall boundaries using cell nuclei as reference (Fig. 4A_1_, A_2_). The flattening of fixed specimens prevented the use of complete three-dimensional (x-y-z) coordinates to characterize algal distance and distribution around the mouth opening. To address this limitation, we arbitrarily selected a range encompassing 20% of the confocal stacks surrounding the focal plane of the mouth for each juvenile. Algae located outside this range were counted to assess displacement along the z-axis and were considered “far from the mouth opening” (Fig. 2SD). Conversely, the x-y coordinates of algal cells within these layers were used to measure their distance from the mouth. These distance values were further normalized by dividing them by the potential maximum travel distance of the algae, defined as the distance from mouth to tail in the corresponding juvenile.

For adults, images were processed as previously described to extract the number of animal nuclei and algae (Gerst et al., 2023; Pinton et al., 2026). Length (excluding the tail) and width were measured on LSM 980 preview images with Zeiss ZEN lite microscopy software v3.11.

### Histology and Transmission Electron Microscopy

*C. macropyga* adults and juveniles (0-2 days after hatching, exposed to *Tetraselmis* sp. for 4-5 days) were relaxed adding drops of 7% MgCl_2_ to filtered artificial sea water and then fixed with a mixture of 0.4% paraformaldehyde and 2.5% glutaraldehyde in 50 mM cacodylate buffer containing 23 mg/ml NaCl and 2.5 g/ml MgCl_2_. The samples were washed in 50 mM cacodylate buffer with 23 mg/ml NaCl, then post-fixed in 1% OsO_4_ in 0.1 M cacodylate buffer, washed with distilled water, and an increasing series of ethanol concentrations and isopropanol were used for dehydration. After infiltration in a mixture of isopropanol and Spurr resin, the samples were embedded in pure Spurr resin for 1 to 2 days at 60°C. *Tetraselmis* algae from liquid cultures were pelleted by centrifugation (1000g, 5min) before fixing and washing as described above. They were then pelleted by centrifugation (15 000 g for 10 min) and embedded in 2% agarose, before proceeding to postfixation.

A Leica EM UC 7 ultramicrotome (Leica Microsystems, Wetzlar, Germany) was used to prepared semithin (600-800 nm) and ultrathin (70 nm) sections from two *C. macropyga* adults (two transversely, one longitudinally) and 5 juveniles (4 transversely, one longitudinally). Semithin sections were stained with a mixture of methylene blue and toluidine blue, mounted with Pertex (MEDITE GmbH, Burgdorf, Germany), and imaged using coherence contrast on an AxioObserver 7 equipped with Axiocam 305 mono camera (Carl Zeiss Microscopy GmbH, Jena, Germany). Images of a total of 14 sections from the 3 adult samples were acquired and stitched with ZEN software (Carl Zeiss Microscopy GmbH, Jena, Germany); damaged, intact, and uncertain algal cells were manually annotated and automatically counted with QuPath v0.5.1 (Bankhead et al., 2017). Ultrathin sections, mounted on slot grids, were contrasted with UranyLess EM Stain (Electron Microscopy Sciences, Hatfield, PA, USA) and 3% lead citrate for 3 min and 2 min, respectively. The samples were examined with a Tecnai 12 transmission electron microscope (FEI Deutschland GmbH, Dreieich, Germany), equipped with a digital camera (TEMCAM FX416, TVIPS, Gauting, Germany).

### Differential gene expression analysis

Bulk RNA extraction, sequencing and transcriptome assembly were carried out as previously described from ca. 100 juveniles 0-3 days post hatching not exposed to algae (5 biological replicates) and 5-10 adults (4 replicates), together with RNA from adults exposed to a bacterial pathogen (Pinton et al., 2026). Trinity isoforms were collapsed to supertranscripts (Davidson et al., 2017) with the script Trinity_gene_splice_modeler.py from Trinity software v2.15.1 (Grabherr et al., 2011), before predicting open reading frames and coding sequences for each supertranscript with TransDecoder v5.5.0 (Haas, 2018) and performing functional annotation with Trinotate v4.0.2 (Altschul et al., 1990; Ashburner et al., 2000; Bryant et al., 2017; Buchfink et al., 2021; Cantalapiedra et al., 2021; Kanehisa et al., 2012; Potter et al., 2018; Powell et al., 2012; The Gene Ontology Consortium et al., 2023). We used bowtie v2.5.4 (Langmead & Salzberg, 2012) to index the transcriptome and align reads, samtools (Danecek et al., 2021) to convert bam to sam files, sort and index them, then UMI-tools *dedup* (T. Smith et al., 2017) to deduplicate the reads. After randomisation with samtools *collate*, the deduplicated reads were quantified with salmon v0.13.1 (Patro et al., 2017) in alignment-based mode. Transcript level estimates, imported into R v4.3.0 (R Core Team, 2023) with the *tximport* package (Love et al., 2018), were analysed with the *DESeq2* package (Soneson et al., 2015): principal component analysis after variance stabilizing transformation was performed with *plotPCA* and differential gene expression analysis with *DESeq*. Gene Set Enrichment Analysis (GSEA) was performed with fgsea within ClusterProfiler (Korotkevich et al., 2016; Wu et al., 2021) on genes ranked by log2fC*(-log10 p) (Xiao et al., 2014), using GO (Gene Ontology) and KEGG (Kyoto Encyclopedia of Genes and Genomes) terms from EggNOG Mapper withing the Trinotate annotation (columns EggNM.GOs, EggNM.KEGG_Module). We referred to sprot_Top_BLASTX_hit of Trinotate annotation to understand the clade the core enrichment genes belong to.

### Statistical analyses and figures

Statistical analyses were carried out in R v4.3.0 (R Core Team, 2023). When needed, a transformation was used (logarithm, square root) to meet model assumptions, according to the Box Cox transformation and Akaike’s information criterion (Akaike, 1998; Crawley, 2013).

Symbiont counts per region (parenchyma, body wall, ventral body wall, dorsal body wall, flanks) were obtained from QuPath, together with their status (healthy, damaged, uncertain) and region areas. Symbiont density (number of cells/area) in body wall and parenchyma of a given image were considered as paired samples and compared with a two-sided Wilcoxon signed rank exact test (*wilcox.test*), given the non-normal distribution of the data. Symbiont densities in the ventral and dorsal sides of the body wall were analysed the same way. To compare the proportion of damaged symbionts among regions (parenchyma, dorsal body wall, ventral body wall, flanks), we excluded algae of unknown status; the proportion was computed with cbind (number of damaged algae vs number of healthy algae) and fitted to a generalised linear mixed model (*glmer* package) with a binomial distribution function (Crawley, 2013). The region was considered as fixed explanatory variable and sample (*i.e.* individual) and slide (*i.e.* image) as nested random explanatory variables for the intercept. Overdispersion and zero inflation were tested for and not detected. The maximal model cbind(Num.Damaged_algae,Num.Healthy_algae) ∼ region + (1|sample/slide) was simplified by removing factors one by one and keeping each simplified model only if equivalent to the previous more complex one (anova, α = 0.05) (Crawley, 2013).

Variation in the ratio of juveniles containing algae at different exposure times was tested with a Pearson’s Chi-squared test. The logarithm of the number of algae within juveniles was fitted to a linear model, with exposure time, PED, or DES as the explanatory variable. The model assumption were tested and a comparison to the null model was used to find the minimal adequate model (Crawley, 2013). Pearson’s chi-squared tests were used to assess differences across timepoints in ratios of algae displaced from the mouth, ratios of juveniles with algae in the body wall, and ratios of algae within the body wall. The distance of algae from the mouth of juveniles, either as a square root of the absolute value or as a ratio of the maximum travelling distance within the animal, was fitted to a linear model with DES as explanatory variable. The significance of the model was determined by comparing it to the null model.

The direction of algal displacement from the juvenile mouth was fitted to a generalised linear mixed model (*glmer* package) with a binomial distribution function as a proportion computed with cbind (dorsal vs ventral) (Crawley, 2013). In order to obtain the intercept value, no fixed effect explanatory variable was given; exposure time and PED were considered as nested random explanatory variables for the intercept, to exclude any possible confounding effect.

For adults exposed to *Tetraselmis* algae, artemia, or FASW (controls), we compared the logarithm of the ratio between algal cells and animal cells and animal size (width * length) across treatments. Student’s t-tests with Bonferroni correction were used to perform pairwise comparisons.

Plots were created with *ggplot2* (Wickham, 2016) and the tree with *ggtree* (Yu et al., 2017) in R v4.3.0 (R Core Team, 2023). Schemes were drawn in Adobe Illustrator, which was also used to annotate the phylogenetic tree; figure plates were assembled with Adobe Indesign. When needed, contrast or brightness were adjusted on the whole picture in Adobe Photoshop or Fiji (Schindelin et al., 2012).

## Data availability

The transcriptomic data are available at NCBI SRA database (BioProject PRJNA1332086, SRAs SRX30605405, SRX30605406, SRX30605407, SRX30605408, SRX30605409, SRX30605410, SRX30605411, SRX30605416, SRX30605417).

## Acknowledgments

We thank all current and former members of the Hejnol working groups at the Friedrich Schiller University Jena and the University of Bergen for their support. We thank Sandor Nietzsche and the Centre for Electron Microscopy of Jena University Hospital - Friedrich Schiller University Jena, as well as Prof. Dr. Elisabeth Liebler-Tenorio and the Institute of Molecular Pathogenesis – Friedrich Loeffler Institute in Jena for access and support with transmission electron microscopy. We thank Dr. Marco Groth and the Core Facility Next-Generation Sequencing of the Leibniz Institute on Aging, Fritz Lipmann Institute in Jena for their support with Illumina sequencing. We thank the University Computer Centre of the Friedrich Schiller University Jena for providing access to the HPC cluster “Draco” used for transcriptomic data analysis.

