## Supplementary material for "Photosymbiotic algae acquisition and their interactions with the acoel *Convolutriloba macropyga*": Supplementary.docx

**Supplementary Figures**

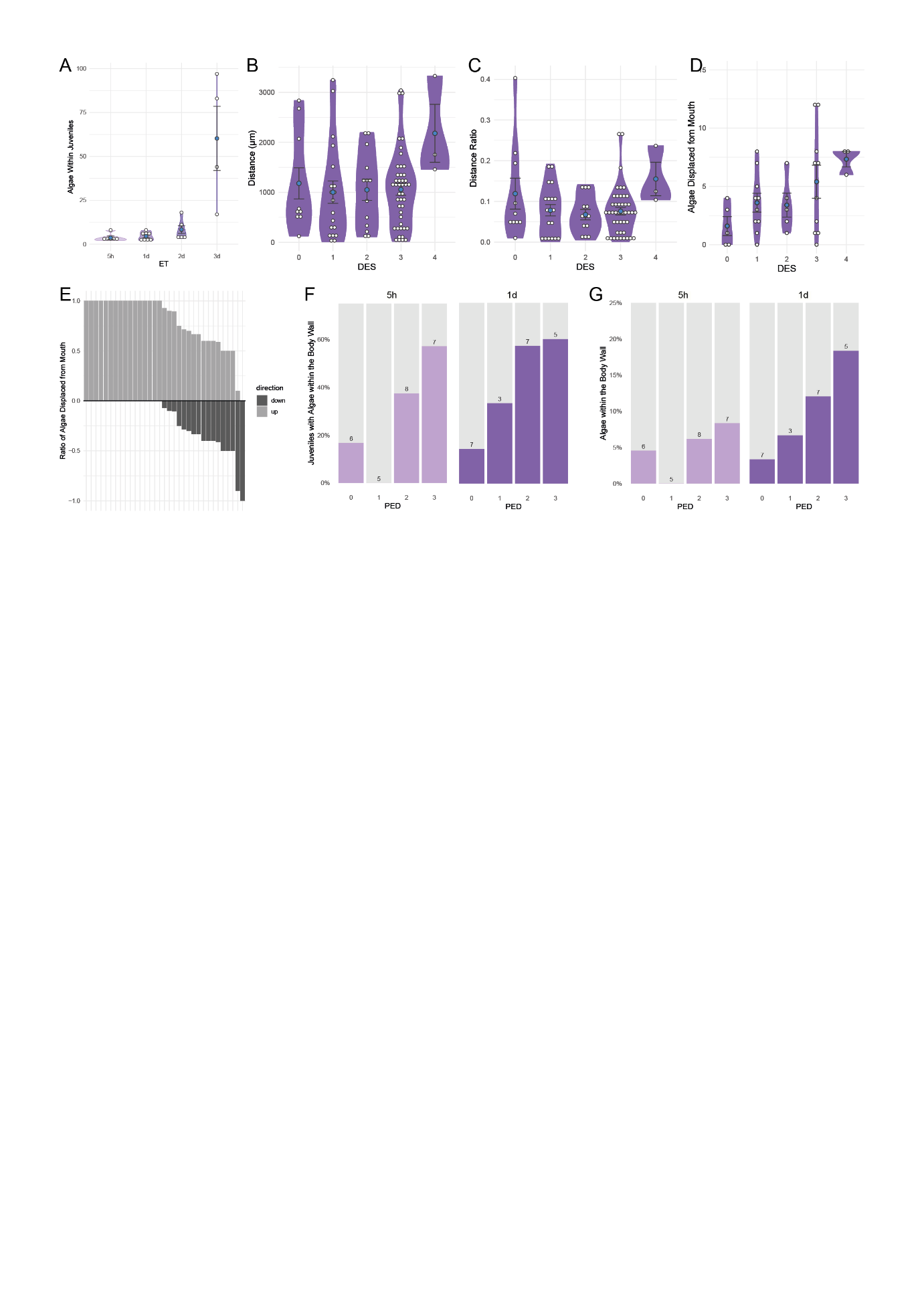

**Fig. S1 Uptake of *Tetraselmis* sp. by juveniles**

**A** Number of algae within *C. macropyga* juveniles at PED 0 (minimal adequate model: log(Algae_number) ~ ET, R^2^= 0.704, p= 4.083e-06). **B** Distance of algae from mouth opening (maximal model: sqrt(distance) ~ DES, minimal adequate model: sqrt(distance) ~ 1). **C** Distance of algae from mouth opening normalized to juveniles’ body size (maximal model: ratio_distance ~ DES, minimal adequate model: ratio_distance ~ 1). **D** Number of algae far from the mouth relative to z axis (minimal adequate model: log(algae_num) ~ DES, R^2^= 0.104, p= 0.133). **E** Ratio of algae outside of the mouth region, by side (cbind(Dorsal, Ventral) ~ 1 + (1 | ET/PED), fixed effects: intercept estimate= 1.3130, z=7.905, p= 2.67e-15; random effects: SD=0, Var=0 for both PED:ET and ET). **F** Percentage of juveniles with algae associated with the body wall (5h ET: χ^2^= 5.238, p= 0.155; 1d ET: χ^2^= 3.640, p = 0.303); sample size above columns. **G** Number of algae associated with the body wall (Pearson’s Chi-squared tests, 5h ET: χ^2^= 3.054, p= 0.383; 1d ET: χ^2^= 5.461, p= 0.141); sample size above columns.

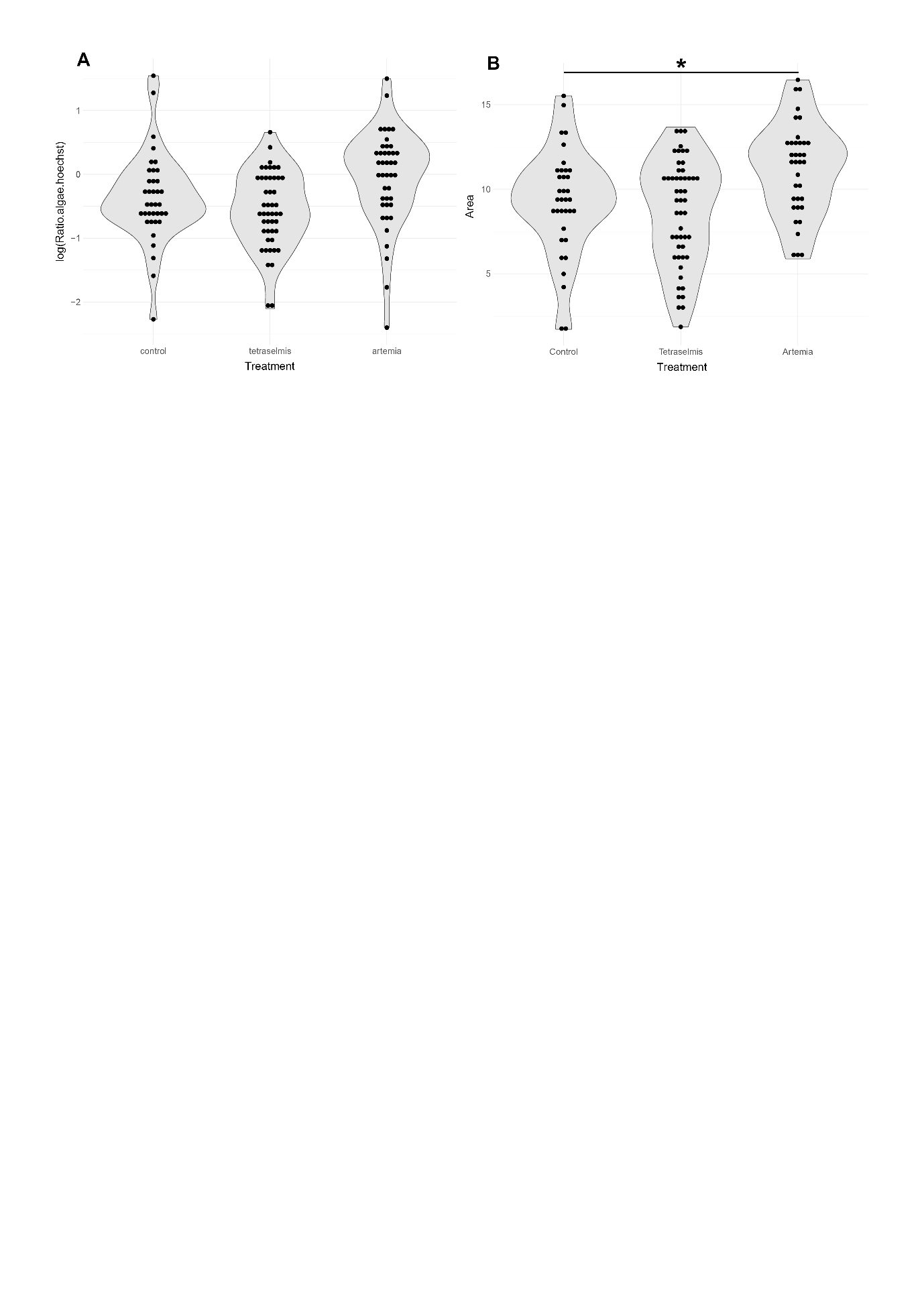

**Fig. S2 *C. macropyga* adults exposed to *Tetraselmis* and *Artemia***

Violin plots of measurements in adult *C. macropyga* exposed to filtered artificial sea water (Control), free-living potential symbionts (*Tetraselmis*), or food (*Artemia*): **A** logarithmic ratio of algal cells to animal cells. Pairwise comparisons: t tests with Bonferroni adjustment for multiple tests: Control-*Tetraselmis*: t = 1.3516, p = 0.3616; Control-*Artemia*: t = -1.988, p = 0.09852. **B** Length x width of individuals. Pairwise comparisons: t tests with Bonferroni adjustment for multiple tests: Control-*Tetraselmis*: t = 0.88487, p = 0.7582; Control-Artemia: t = -2.8052, p = 0.013142. *: p < 0.05

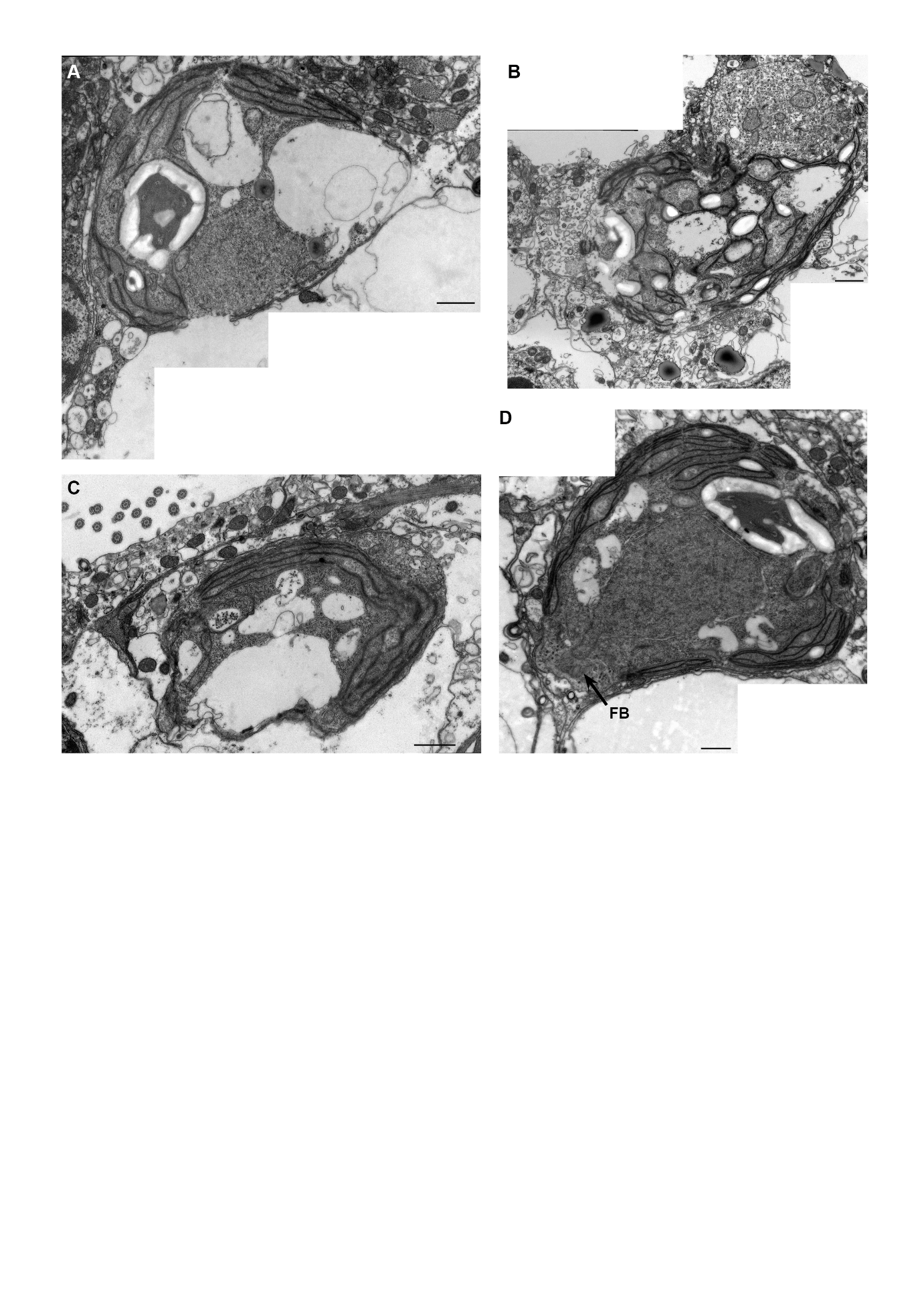

**Fig. S3 Further pictures of algae in juveniles, including damaged ones**

FB: Flagellar Bases. Scale bars: 1µm.

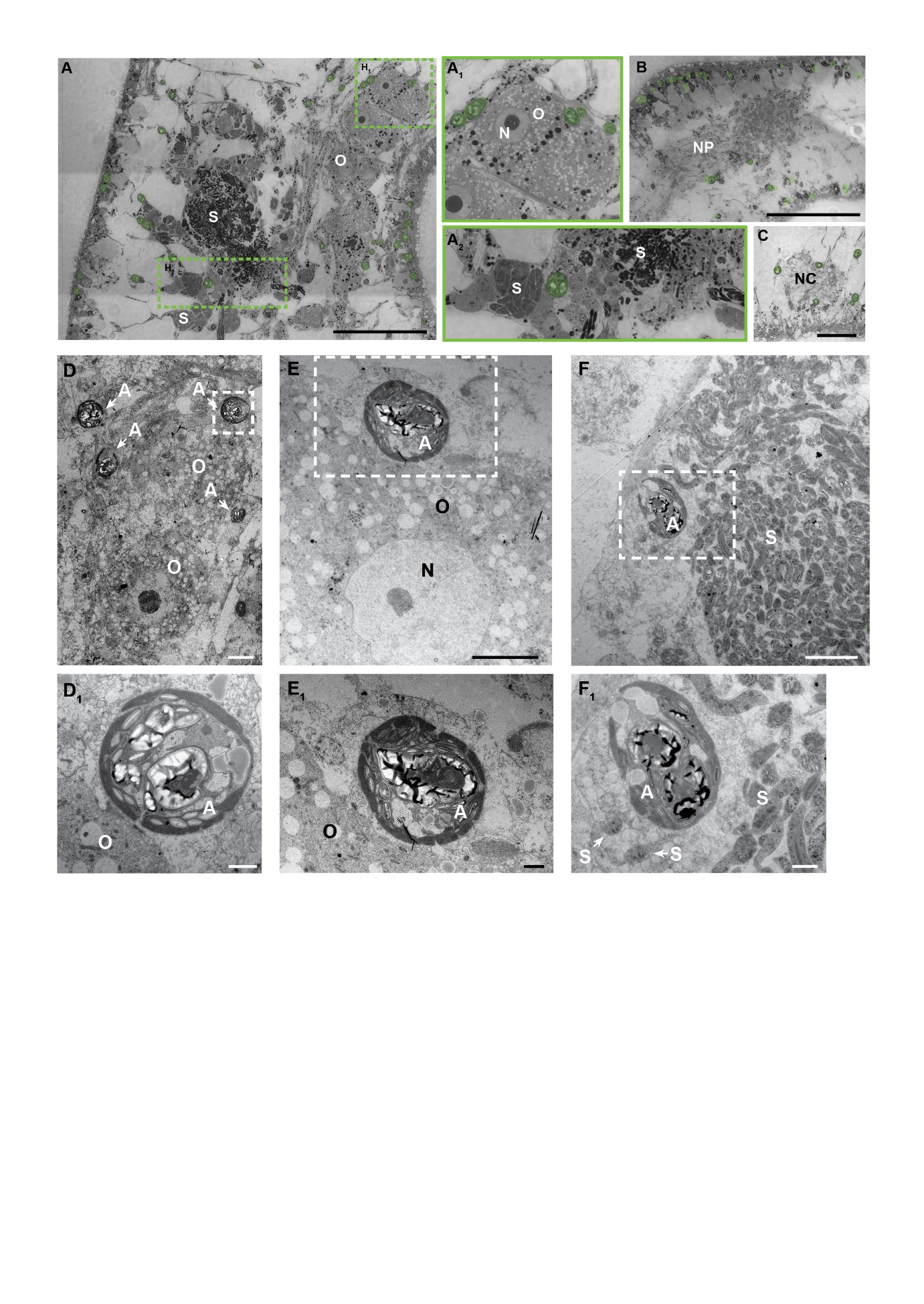

**Fig. S4 TEM pictures of symbionts close to *C. macropyga* gonads and nervous system**

Histological cross sections with algal symbionts pseudocoloured in green: **A** at the level of the gonads (dorsal side facing left); **A_1_, A_2_** higher magnification of the dashed rectangles in **A**. **B** at the level of the anterior nervous system; **C** showing a nerve cord. in **A, B, C**. TEM photographs of algal symbionts in close association with oocytes (**D,E** – details in **D_1_,E_1_**) and sperm (**F** – detail in **F_1_**). A: algal symbionts, N: nucleus of the oocyte, NC: nerve cord, NP: neuropil, O: oocyte; S: sperm cells. Scale bars are 5 µm in **A, B, C**, 10 µm in **D, E, F** and 2 µm in **D_1_,E_1_,F_1_**.

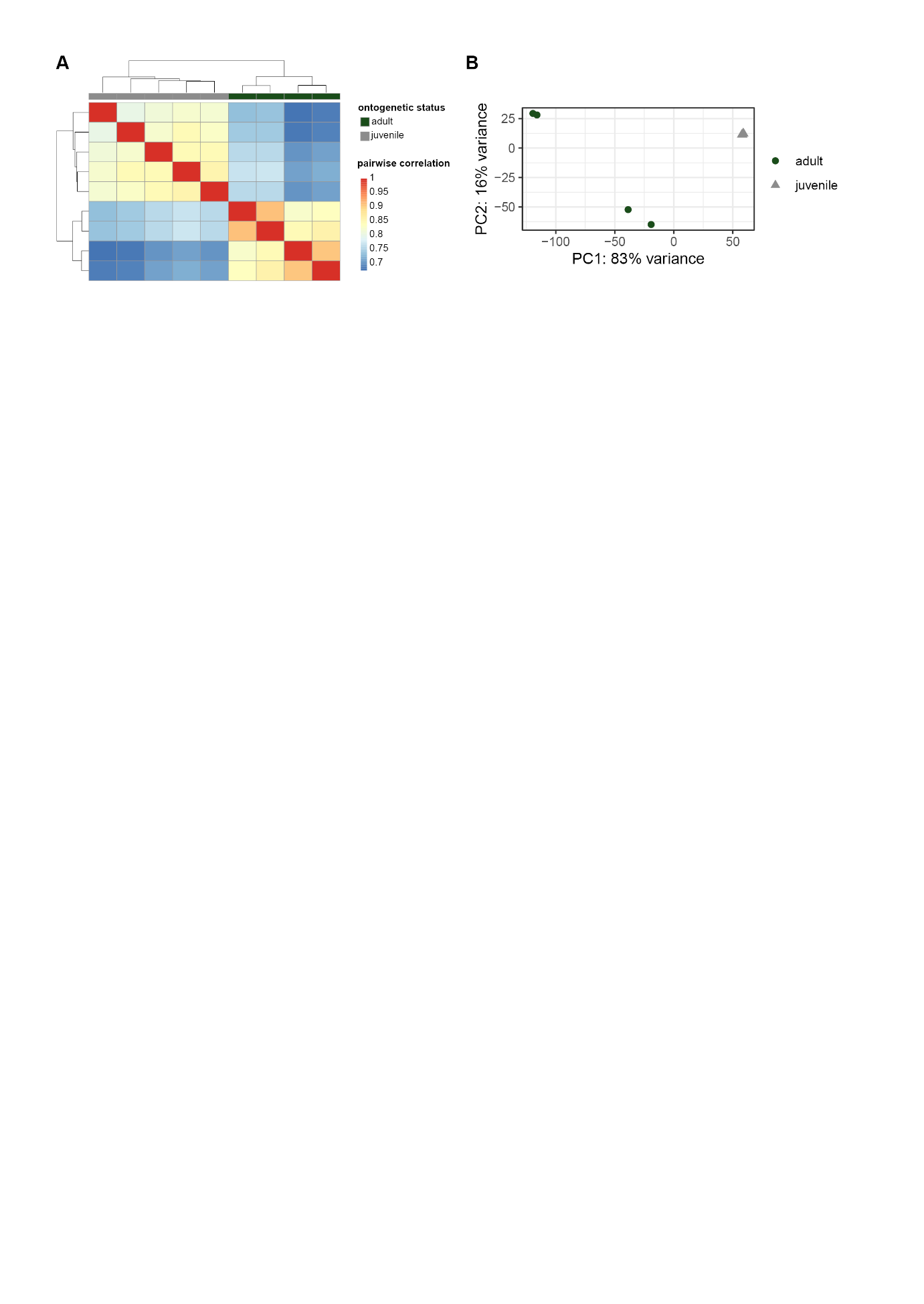

**Fig. S5 transcriptomic dataset analysis**

Analysis of DESeqDataSet after variance stabilizing transformation. **A** heatmap of the pairwise correlation value. **B** PCA plot.

**Supplementary Data**

**Video S1, S2, S3 juveniles feeding on *Tetraselmis***

**Video S4 *Tetraselmis* sp. SAG 35.93 in culture**

**Data S1 rbcL sequences in fasta format**

**Data S2 rbcL phylogeny in Newick format**

**Table S1 GSEA GO terms results (csv)**

**Table S2 core enrichment genes of GSEA GO terms (csv)**

**Tables S3 GSEA KEGG results (csv)**

**Table S4 core enrichment genes of GSEA KEGG (csv)**
